# Distinct small molecule inhibitors of Kras specifically prime CTLA4 blockade therapy to transcriptionally reprogram Tregs and overcome resistance to suppress pancreas cancer

**DOI:** 10.1101/2025.02.28.640711

**Authors:** Krishnan K. Mahadevan, Ana S. Maldonado, Bingrui Li, Aaron A. Bickert, Adrian Kapercyzk-Perdyan, Shreyasee V. Kumbhar, Sujan Piya, Amari Sockwell, Sami J. Morse, Kent Arian, Hikaru Sugimoto, Shabnam Shalapour, David S. Hong, Timothy P. Heffernan, Anirban Maitra, Raghu Kalluri

## Abstract

Lack of sustained response to oncogenic Kras (Kras*) inhibition in preclinical models and patients with pancreatic ductal adenocarcinoma (PDAC) emphasizes the need to identify impactful synergistic combination therapies to achieve robust clinical benefit. Kras* targeting results in an influx of T cell infiltrates including Tregs, effector CD8^+^ T cells and exhausted CD8^+^ T cells expressing several immune checkpoint molecules in PDAC. Here, we probe whether the T cell influx induced by different Kras* inhibitors enable a therapeutic window to prime adaptive immune response in PDAC. Here we report a specific synergy between Kras^G12D^ allele specific inhibitor, MRTX1133 or multi-selective pan-RAS inhibitor, RMC-6236 and anti-CTLA4 immune checkpoint blockade. In contrast, attempted therapeutic combination with multiple other immune checkpoint inhibitors, including anti-PD1, anti-Tim3, anti-Lag3, anti-Vista and anti-4-1BB agonist antibody failed due to compensatory mechanisms mediated by other checkpoints on exhausted CD8^+^ T cells. Specifically, anti-CTLA4 therapy in Kras* targeted PDAC transcriptionally reprograms effector T regs to a naïve phenotype, reverses CD8^+^ T cell exhaustion and is associated with recruitment of tertiary lymphoid structures (TLS) containing follicular B cells, interferon (IFN)- stimulated/ activated B cells, plasma cells and germinal center B cells to functionally enable efficacy of immunotherapy with long-term survival. In this regard, inhibition of the TLS with lymphotoxin-β inhibitor (LTBi) or direct B cell depletion reversed the survival benefit conferred by the combination therapy and highlights the function of TLS in generating productive anti-tumor immune responses. Further, single cell ATAC sequencing analysis revealed that transcriptional reprogramming of Tregs is epigenetically regulated by downregulation of AP-1 family of transcription factors including Fos, Fos-b, Jun-b, Jun-d in the IL-35 promoter region. This study reveals an actionable vulnerability in the adaptive immune response in Kras* targeted PDAC with relevant clinical implications.

## Introduction

Pharmacological targeting of oncogenic Kras (Kras*), a mutation that was considered ‘undruggable’ until recently represents a significant breakthrough in the treatment of several malignancies and has the potential to benefit numerous cancer patients. Strategies to inhibit oncogenic Kras include small molecule inhibitors of Kras^G12D^ (MRTX1133, RMC-9805, ASP3082 and KRB-456) ^1–5^ and Kras^G12C^ mutations (Sotorasib/ AMG-510^6–8^, Adagrasib/ MRTX849 ^9–11^), pan-Ras inhibitors (RMC-6236, RMC-7977) ^12,13^, pan Kras inhibitors (BI-2493, BI-2865) ^14^, iExosomes inhibiting Kras^G12D^ ^15,16^and drugs inhibiting downstream targets of oncogenic Kras such as MEK and ERK ^17,18^. Among the different oncogenic Kras mutations, G12C mutations are frequent in non-small cell lung cancers (NSCLCs) and colorectal cancers, whereas G12D mutations are frequent in PDAC and cholangiocarcinoma ^19–21^. Other frequently mutated Kras genes include G12R and G12V mutations in PDAC ^22^. Kras^G12D^ represents the most frequently mutated gene in pancreatic ductal adenocarcinoma (PDAC) ^22,23^.

Targeting Kras^G12D^ with small molecule inhibitors such as MRTX1133, BI-2493, RMC-6236, and iExosomes resulted in robust increase in survival in mouse models of PDAC ^1,3,7,10,14–16,24^. However, prolonged treatment with small molecule inhibitors of Kras* eventually resulted in development of resistance to Kras* targeting and tumor relapse in preclinical models of PDAC^18,24–27^. Further, Sotorasib and Adagrasib (G12C inhibitors) have resulted in close to 30-50% response rates in non-small cell lung cancers (NSCLCs), pancreatic and biliary tract carcinomas but disease progression/ relapse was observed in clinical trials ^11,28–30^. Several strategies to address cancer cell intrinsic mechanisms of resistance to genetic or pharmacological Kras* suppression include targeting CDK4/6, YAP, CDK8, TEAD, MYC and SOS, in addition to targeting EMT, proteostasis networks and EGFR eventually lead to relapse despite longer survival outcomes in preclinical models of PDAC ^18,25–27,31–37^. Therefore, identification and leveraging tumor microenvironment dependent vulnerabilities resulting from Kras* targeting is critical to achieve meaningful treatment outcomes in PDAC patients.

Prior studies with genetically engineered mouse models using an inducible Kras* allele demonstrated that Kras* is a dominant driver of T cell suppression in colorectal cancers and PDAC^38–40^. Kras* targeting across different cancer models led to a global increase in T cell infiltrates including CD4^+^ T cells, Tregs and CD8^+^ T cells in the tumor microenvironment (TME) indicating that the downstream effects of Kras* targeting could result in targetable vulnerabilities in the adaptive immune response ^38–46^. Kras* targeting has shown divergent vulnerabilities in the immune response across cancer models in an organ specific context ^38,41,44,46^. Therefore, we hypothesized that the immune microenvironment remodeling following Kras* targeting may offer new vulnerabilities critical to inform specific combination therapies in PDAC.

In this study, we examine the adaptive immune response following targeting Kras^G12D^ with MRTX1133 and pan-RAS inhibitor, RMC-6236 in PDAC. Our study reveals that despite the use of distinct Kras* targeting strategies, similar recruitment of T cells in pre-clinical mouse models was observed. We identify a unique synergy between Kras* targeting and anti-CTLA4 mediated by transcriptional regulation of Tregs and enhanced CD8^+^ T cell responses in conjunction with recruitment of tertiary lymphoid structures (TLS), all associated with suppression of PDAC and prolonged survival. This study informs rational combination strategies for clinical testing.

## Results

### Pharmacological inhibition of Kras* results in parallel recruitment of effector and suppressor T cells in pre-clinical mouse models

To assess alterations in the PDAC TME following Kras* targeting, we performed single cell RNA sequencing (scRNA seq.) analysis in orthotopic KPC1 PDAC (primary PDAC line derived from 343P KPC-*Pdx1-Cre; LSL-Kras^G12D/+^; Trp53^R172H/+^*; *Rosa-LSL-EYFP* tumor bearing mice) treated with Kras^G12D^ specific small molecule inhibitor, MRTX1133 (**Fig. 1A**). MRTX1133 treatment was started once the KPC1 mice developed palpable, large PDAC (**Fig. 1A**). UMAP analysis of the scRNA seq. captured fibroblasts, cancer/ ductal cells, acinar cells, T cells, innate lymphoid cells (ILCs), endothelial cells, dendritic cells (DCs), myeloid cells, B−/plasma cells and granulocytes in the PDAC tissue (**Supplementary Fig. S1A-C**). MRTX1133 treatment led to a significant increase in the proportion of T and B cells, accompanied by a decrease in myeloid cells, cancer/ ductal cells and fibroblasts ^40,41^ (**Fig. 1B-E, Supplementary Fig. S1A, B**).

**Figure 1:**
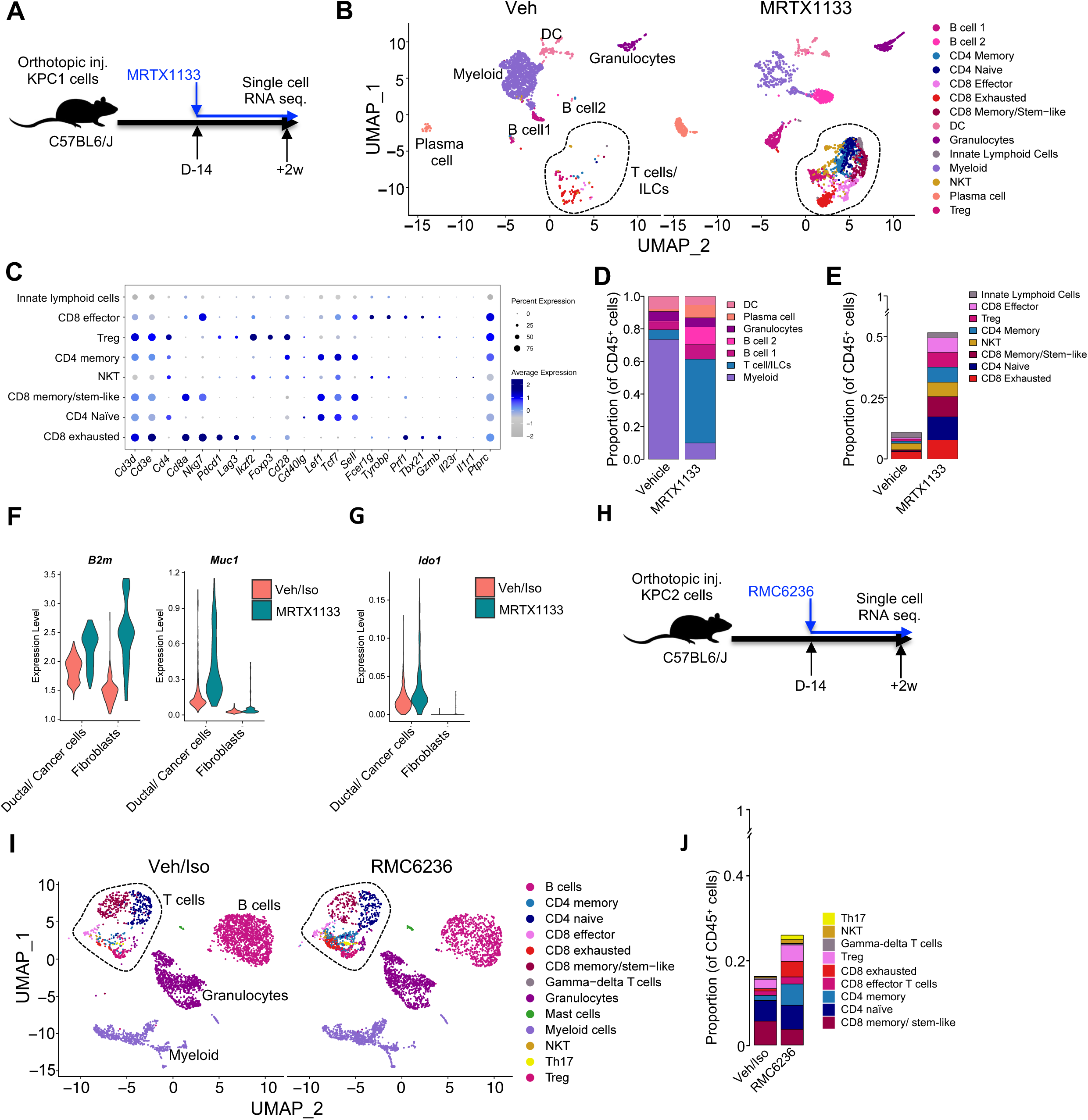
Kras* targeting led to a global increase in T cell infiltrates in PDAC. **(A)** Schematic representation of orthotopic injection and MRTX1133 treatment of KPC1 mice. **(B)** UMAP of all cell types with T cell subsets in Vehicle/ MRTX1133 treated KPC1 PDAC determined by scRNA seq. analysis (n=3 tumors/ group). **(C)** Dot plots of genes used to define T cell subtypes in (B). **(D-E)** Relative abundance of all cell types (D) and T cells as a percentage of CD45^+^ cells (E) in KPC1 PDAC determined by scRNA seq. **(F-G)** Violin plots for *B2m*, *Muc1* **(F)** and *Ido1* **(G)** in ductal/ cancer cells and fibroblasts of KPC1 mice determined by scRNA seq. (n=3 tumors/ group). **(H)** Schematic representation of orthotopic injection and RMC6236 treatment of KPC2 mice. **(I)** UMAP of all cell types with T cell subsets in Veh/Iso RMC6236 treated KPC2 PDAC determined by scRNA seq. analysis (n=3 tumors/ group). **(J)** Relative abundance of T cells as a percentage of CD45^+^ cells in KPC2 PDAC determined by scRNA seq. Data are presented as mean in **D, E and J** and as violin plots in **F, G**. See related figs. S1.

Analysis of tumor infiltrating T cells revealed a global increase in the number of CD4^+^, CD8^+^ T cells, NK T cells and innate lymphoid cells (**Fig. 1B-D, Supplementary Fig. S1D**). Further sub-clustering of the T cells demonstrated an increase in number and frequencies of CD8 effector cells (*Prf1*, *Tbx21*, *Ccl5*, *Gzmb* high and *Pdcd1*, *Lag3* low), exhausted CD8 cells (*Pdcd1*, *Lag3* high), T regs (CD4 cells with *Ikzf2*, *Foxp3* high and *Tbx21* low), CD4 naïve (*Lef1*, *Tcf7*, and *Sell*) and memory cells (*Cd28*, and *Cd40lg*) (**Fig. 1B-E, Supplementary Fig. S1D-E**). Analysis of cancer/ductal cells and fibroblast populations demonstrated an increase in interferon responsive gene, *B2m* (**Fig. 1F**), a subunit of MHC-1 which aids in the recruitment of CD8^+^ T cell response in PDAC and colorectal cancers (CRCs) ^38,47^. Furthermore, Kras* targeting resulted in upregulation of *Muc1* and its downstream mediator *Ido1* in the cancer cells and fibroblasts which are primary meditators of Treg recruitment (**Fig. 1G**)^48^. Consistent with an increase in Treg recruitment following MRTX1133 treatment, we observed suppression of *Il6* and *Tnf* in the cancer cells and fibroblasts (**Supplementary Fig. S1F**). Analyses of the cytokines involved in T reg recruitment demonstrated an increase in Nfkb1 (NF-κB) and Cxcr*4*/*Ackr3* (Cxcr7) in cancer cells and fibroblasts following Kras* targeting (**Supplementary Fig. S1G**) ^49^.

Next, we use a second orthotopic KPC2 (primary PDAC line derived from HY19636-PKC-*P48-Cre, LSL-Kras^G12D/+^, Trp53^L/+^* mice) tumor bearing mice with the pan-RAS inhibitor, RMC6236 to determine whether the changes in T cell infiltrates following Kras* inhibition are conserved across multiple PDAC models and inhibitors (**Fig. 1H**). We sorted for CD45^+^ lymphocytes to enrich for the immune population in the TME. UMAP analysis of the scRNA seq identified T cells, B cells, granulocytes, mast cells and myeloid cells (**Fig. 1I**). Analysis of the tumor infiltrating T cells revealed an increase in the frequencies of CD4 memory, CD4 naïve cells, T regs, effector and exhausted CD8^+^ T cells (**Fig. 1I-J**). Taken together, our results indicate that Kras* targeting resulted in an influx of effector and suppressor T cell infiltrates into the TME in two PDAC models with different pharmacological inhibitors.

### Kras* targeting synergizes with anti-CTLA4 but not anti-PD1 therapy

The recruitment of distinct T cell subtypes following Kras* inhibition presented an opportunity to target therapeutic vulnerabilities in the adaptive immune responses. Although checkpoint immunotherapy conferred survival benefit in cancers such as melanoma, lymphomas and subsets of lung cancers, PDAC remains refractory to T cell targeted therapies in preclinical models and clinical trials ^50–53^. First, we assessed the efficacy of anti-CTLA4 and anti-PD1 treatment in combination with MRTX1133 and RMC-6236 in KPC2 tumor bearing mice (**Fig. 2A**). Because cancer cells in the KPC1 tumors express EYFP which could result in potential immunogenicity, we use the KPC2 mouse model for subsequent experiments to avoid the fluorescent reporter confounding therapeutic immune responses (**Supplementary Fig. S2A-B**)^54^. MRTX1133 and RMC-6236 significantly prolonged survival and inhibited PDAC progression (**Fig. 2B-K**). Further, analysis for CK19 (a marker of ductal/ cancer cells to assess tumor burden) confirmed significant tumor inhibition upon MRTX-1133 and RMC-6236 treatments (**Supplementary Fig. S3A-I**). Analysis of pERK showed downregulation of pERK demonstrating similar levels of target engagement with MRTX1133 and RMC-6236 (**Supplementary Fig. S3A-F)**. Despite initial suppression of PDAC growth and prolonged survival, MRTX1133 and RMC-6236 treated mice eventually succumbed to PDAC, suggesting resistance to Kras* targeting (**Fig. 2B-C, Supplementary Fig. 3G-J**). Addition of anti-CTLA4 immunotherapy to MRTX1133 and RMC-6236 demonstrated prolonged tumor growth inhibition, survival and in the timeframe studied, the combination prevented the emergence of resistance to Kras* targeting (**Fig. 2B-K, Supplementary Fig. 3A-J)**. In contrast, addition of anti-PD1 with Kras* targeting did not result in an improvement when compared to Kras* targeting alone (**Fig. 2B-K, Supplementary Fig. S3A-J**). Analysis of the immune infiltrates demonstrated an increase in CD8^+^ T cells in MRTX1133 or RMC-6236 treated tumors compared to controls. Addition of anti-CTLA4 further expanded CD8^+^ and CD19^+^ cells in the pancreatic TME likely resulting in sustained PDAC inhibition response following Kras* targeting (**Fig. 2D-K**).

**Figure 2:**
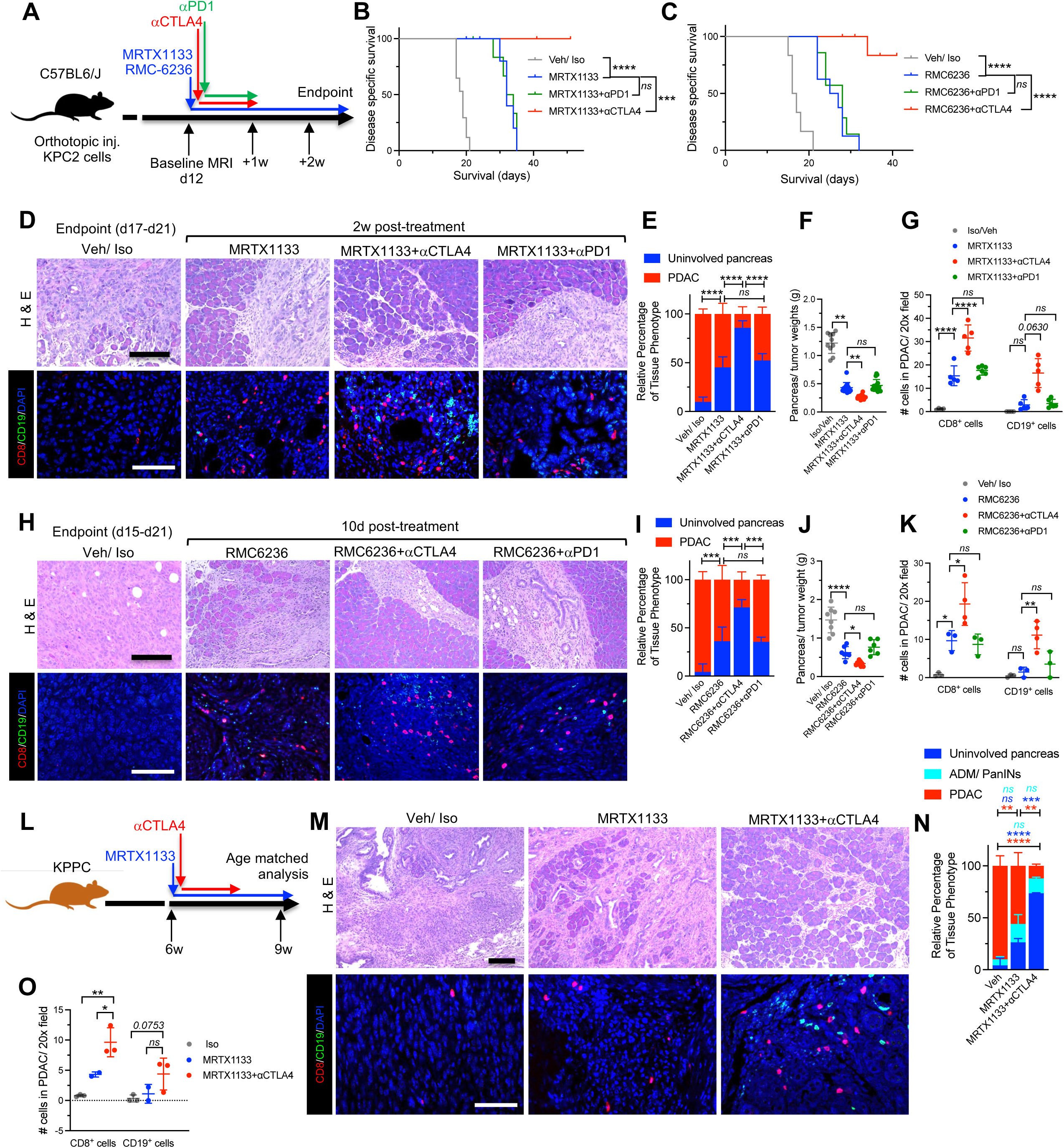
Kras* targeting increased T cell infiltrates and synergized with anti-CTLA4 therapy in PDAC. **(A)** Schematic representation of orthotopic injection and RMC6236, MRTX1133, aCTLA4 and aPD1 treatments of KPC2 mice. **(B-C)** Kaplan-Meier survival curves of Veh/Iso (n=17), MRTX1133 (n=11), MRTX1133+aCTLA4 (n=11), MRTX1133+aPD1 (n=9) treated KPC2 mice **(B)**, Veh/Iso (n=6), RMC6236 (n=8), RMC6236+aCTLA4 (n=10), RMC6236+aPD1 (n=8) treated KPC2 mice **(C)**. **(D-E)** Representative H&E (top panel) and CD8-CD19-DAPI immunofluorescence images (bottom panel) from MRTX1133 treated age-matched KPC2 mice with aCTLA4 or aPD1 **(D)**, with quantification of tissue histology **(E)**. Veh/Iso (n=3), MRTX1133 (n=11), MRTX1133+aCTLA4 (n=7), MRTX1133+aPD1 (n=7). **(F)** Pancreas/ tumor weights of age-matched KPC2 mice treated with Veh/Iso (n=10), MRTX1133 (n=13), MRTX1133+aCTLA4 (n=15), MRTX1133+aPD1 (n=15). **(G)** Quantification of CD8 and CD19^+^ cells from **D** (n=4-6 mice/group). **(H-I)** Representative H&E (top panel) and CD8-CD19-DAPI immunofluorescence images (bottom panel) from RMC6236 treated age-matched KPC2 mice with aCTLA4 or aPD1 **(H)**, with quantification of tissue histology **(I)** (n=3/group). **(J)** Pancreas/ tumor weights of age-matched KPC2 mice treated with Veh/Iso (n=8), RMC6236 (n=7), RMC6236+aCTLA4 (n=9), RMC6236+aPD1 (n=6). **(K)** Quantification of CD8 and CD19^+^ cells from **H** (n=3-4 mice/group). **(L)** Schematic representation of MRTX1133, and aCTLA4 treatments of autochthonous KPPC mice. **(M-O)** Representative H&E (top panel) and CD8-CD19-DAPI immunofluorescence images (bottom panel) from MRTX1133 treated age-matched KPPC mice with aCTLA4 **(M)**, quantification of tissue histology **(N)**, and CD8 and CD19^+^ cells **(O)** from **M** (KPPC Veh/Iso: n=3, MRTX1133: n=2, MRTX1133+aCTLA4: n=3). <u>Note:</u> One Veh/Iso mice in M has been used earlier in another study^41^. Data are presented as mean ± SD in **E, F, G, I, J, K, N** and **O**. Significance was determined by log-rank test in **B** and **C**, two-way ANOVA in **E, I** and **N**, Kruskal-Wallis with Dunn’s multiple comparisons test in **F** and CD19 comparisons in **G**, one-way ANOVA with Dunnett’s multiple comparisons test in **J, K, O** and CD8 comparisons in **G**. Scale bars-100mm. \**P* < 0.05, \*\**P*< 0.01, \*\*\**P* < 0.001, \*\*\*\**P* < 0.0001; *ns*, not significant. See related figs. S2-S4.

We next probed the efficacy of Kras* inhibitors and anti-CTLA4 combination therapy in eliciting robust antitumor responses in autochthonous genetically engineered mouse models (GEMs) (**Fig. 2L**). We treated autochthonous KPPC (*P48-Cre, LSL-Kras^G12D/+^, Trp53^L/L^*) mice with MRTX1133 and anti-CTLA4 therapy following establishment of PDAC starting at 6 weeks of age (**Fig. 2L-N**). Mice were sacrificed for age matched analysis ∼3 weeks after indicated treatment when the vehicle treated mice reached moribundancy (**Fig. 2L-N**). MRTX1133 treated KPPC mice demonstrated significant suppression of PDAC compared to the vehicle treated controls and addition of anti-CTLA4 further resulted in suppression of PDAC with an increase in the proportion of uninvolved pancreatic tissue (**Fig. 2M-N**). Analysis of tumor infiltrating lymphocytes demonstrated an increase in CD8^+^ T cells and an increasing trend in CD19^+^ B cells with MRTX1133 and anti-CTLA4 combination therapy compared to MRTX1133 monotherapy (**Fig. 2M-O**).

Next, we used autochthonous KPC (*Pdx1-Cre, LSL-Kras^G12D/+^, Trp53^R172H/+^)* GEM mice to determine if the combination therapy demonstrates synergy in KPC mice (**Supplementary** Fig. 4A). All vehicle (4 out of 4) treated KPC mice demonstrated invasive PDAC 10 weeks post-treatment (**Supplementary** Fig. 4B-C). Age matched analysis of mice treated with MRTX1133 demonstrated suppression in 3 out of 4 tumors and 1 mouse demonstrated resistance to Kras* inhibition (**Supplementary** Fig. 4B-C). Mice that received combination therapy demonstrated minimal PDAC and suppression in 3 out of 3 mice (**Supplementary** Fig. 4B-C).

To further validate our findings, we used a different KPC3 orthotopic tumor model (primary PDAC line derived from KPC T*-Pdx1-Cre, LSL-Kras^G12D/+^*, Trp53^L/+^ derived cell line) for long-term follow up after treatment with MRTX1133 and anti-CTLA4 (**Supplementary** Fig. 4D-E). Anti-CTLA4 as a monotherapy did not increase survival of PDAC tumor bearing mice, as shown before^52^,whereas the combination of MRTX1133 and anti-CTLA4 demonstrated sustained tumor suppression in majority of the PDAC mice (**Supplementary** Fig. 4D-E). Our results indicate that MRTX1133 and RMC-6236 demonstrated therapeutic synergy with anti-CTLA4 irrespective of the Kras* targeting strategy.

### Kras* targeting in combination with anti-CTLA4 drives Tregs towards a naïve state, attenuates CD8 exhaustion and increases effector CD8^+^ T cells

Single cell RNA seq. analysis of the T cells in the KPC3 mice treated with MRTX1133 and anti-CTLA4 showed a decrease in frequencies and number of exhausted CD8^+^ T cells, expansion of memory/stem like, effector CD8^+^ T cells, naïve and memory CD4^+^ T cells (**Fig. 3A-B, Supplementary Fig. S5A-C**). Analysis of scRNA seq. data in a second, KPC2 model treated with RMC-6236 in combination with anti-CTLA4 therapy validated the increase in total T cell infiltrates, expansion of memory/ stem-like and effector CD8^+^ T cells, naïve and memory CD4^+^ T cells (**Fig. 3C-D, Supplementary Fig. S5D-F**). Notably, the anti-CTLA4 combination with Kras* targeting did not decrease total Treg numbers (**Fig. 3A-D**). We used the 9H10 clone of the anti-CTLA4 antibody that blocks CTLA4 binding to the B7 co-receptor in this study (similar to mechanisms of Ipilimumab and Tremelimumab that do not deplete Tregs^55^). Rather, unsupervised analysis of differentially regulated genes (DRGs) in the tumors that received MRTX1133 and anti-CTLA4 combination therapy revealed a distinct transcriptional profile in Tregs compared to the those in anti-CTLA4 or MRTX1133 treated tumors (**Fig. 3E-F**). The Tregs from MRTX1133 and anti-CTLA4 treated tumors demonstrated downregulation of AP-1 related genes including *Rgs2*, *Fos*, *Fosb, Junb and Jund* (**Fig. 3E-F**). Among these, *Rgs2*, *Fos*, *Fosb* have been shown to be involved in differentiation of Tregs. *Junb* promoted effector T reg development through facilitation of IRF4 and *Jund* protected Tregs from apoptosis^56–58^. Further, we observed a downregulation of *Dyrk1a*, *Dusp4*, *Samsn1* and *Ccr5* in the Tregs of mice that received MRTX1133 and anti-CTLA4 combination therapy (**Fig. 3E-F**). *Dyrk1a* expression has been shown to drive CD4^+^ T cells into Tregs differentiation lineages^56–58^, *Dusp4* is known to stabilize Foxp3 expression in T regs^59^, *Samsn1* and *Ccr5* enabled migration and homing ^60^ (**Fig. 3E-F**). Taken together, our analysis indicates that genes involved in Treg differentiation, effector function, homing and migration were all downregulated in the tumor infiltrating Tregs of mice that received MRTX1133 and anti-CTLA4 combination therapy. Analysis of the Tregs in RMC6236 and anti-CTLA4 treated tumors also revealed downregulation several of the genes that promote effector T reg phenotype such as *Rgs2*, *Fos*, *Fosb*, *Junb*, *Dusp4*, and *Ccr5* validating the results with MRTX1133 and anti-CTLA4 treated tumors (**Fig. 3G, Supplementary Fig. S6A-B**). Further, the Tregs from MRTX1133/ RMC-6236 and anti-CTLA4 treated tumors demonstrated upregulation of *Ifi213* which promotes IFN responsive genes (**Fig. 3E-G, Supplementary Fig. S6A**). Next, we analyzed expression of effector Treg cytokines, IL10, TGF-β1 and IL35 in the Tregs of mice that received Kras* targeting and immunotherapy^61–63^. We identified a decrease in *Il10*, *Tgfb1*, and the IL35 subunit *Ebi3* expression in the Tregs of mice treated with MRTX1133 or RMC6236 in combination with anti-CTLA4 indicating suppression of the effector Treg function (**Fig. 3H-I**). Collectively, our data indicates that Kras* targeting in combination with anti-CTLA4 therapy drives Tregs to a naïve state and inhibits effector T reg pathways critical for suppression of CD8^+^ T cells.

**Figure 3:**
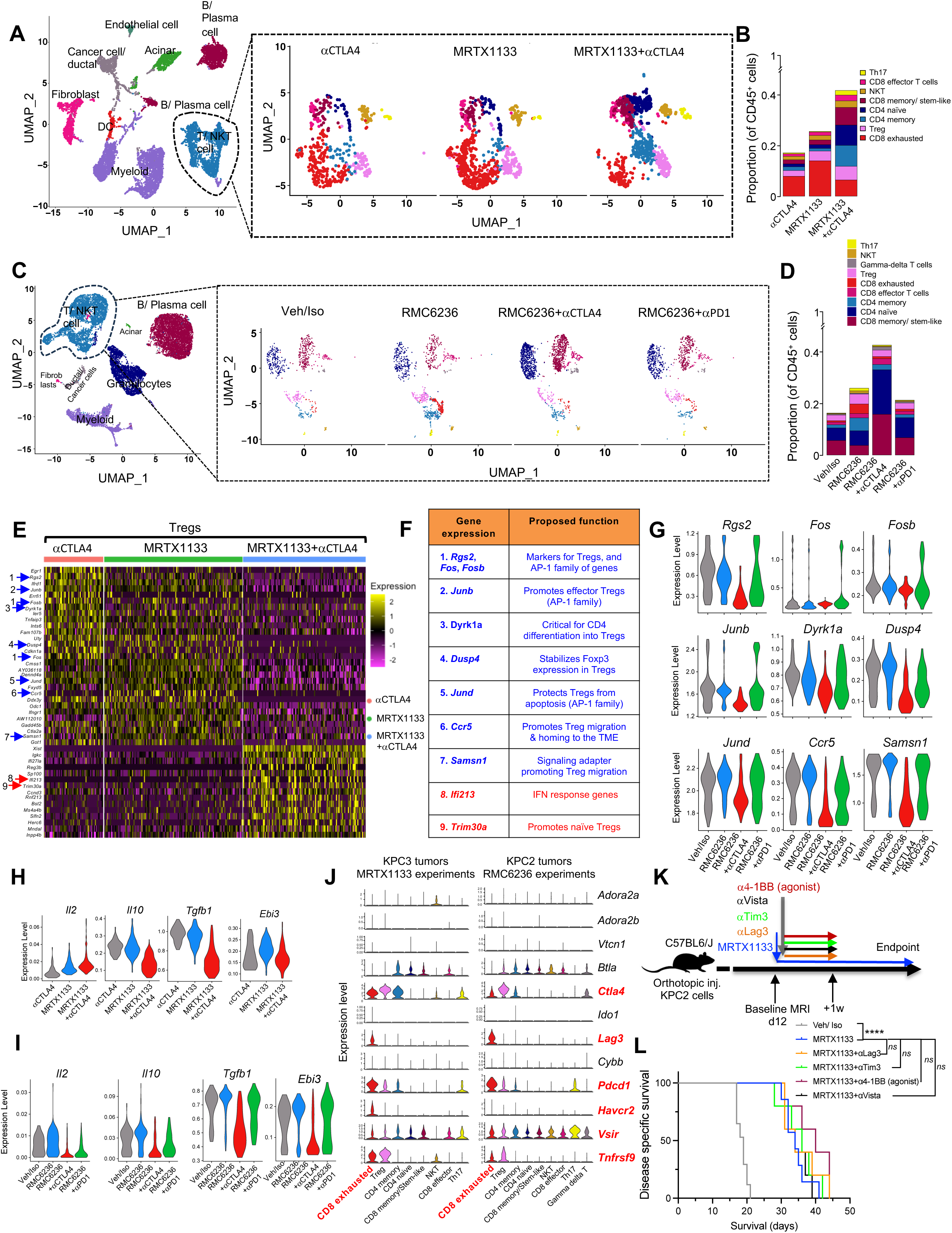
scRNA seq. analysis of T cell infiltrates in MRTX1133, RMC6236 and checkpoint immunotherapy treated PDAC. **(A-B)** UMAP of all cell types (left) and T cell subsets in aCTLA4, MRTX1133, MRTX1133+aCTLA4 treated (right) KPC3 PDAC **(A)**, with relative frequencies of T cell subsets (as % of CD45^+^ cells) **(B)** determined by scRNA seq. analysis (n=3 mice/group). **(C-D)** UMAP of all cell types (left) and T cell subsets in Veh/Iso, RMC6236, RMC6236+aCTLA4, RMC6236+aPD1 treated (right) KPC2 PDAC **(C)**, with relative frequencies (as % of CD45^+^ cells) of T cell subsets **(D)** determined by scRNA seq. analysis (n=3 tumors/group). **(E-F)** Heatmap of differentially regulated genes in the tumor infiltrating Tregs of aCTLA4, MRTX1133, MRTX1133+aCTLA4 treated KPC3 PDAC (Red arrow indicates upregulated genes and blue arrow indicates downregulated genes in the MRTX1133+aCTLA4 group compared to the other groups) **(E)**, with summary table of gene functions **(F)**. **(G)** Violin plots for *Rgs2, Fos, Fosb, Junb, Dyrk1a, Dusp4, Jund, Ccr5* and *Samsn1* in tumor infiltrating Tregs of RMC6236 treated KPC2 mice with aCTLA4/aPD1. **(H-I)** Violin plots for *Il2, Il10, Tgfb1* and *Ebi3* in tumor infiltrating Tregs of MRTX1133 **(H)**, RMC6236 **(I)** treated mice with aCTLA4/aPD1. **(J)** Violin plots for immune checkpoint genes *Adora2a, Adora2b, Vtcn1, Btla, Ctla4, Ido1, Lag3, Cybb, Pdcd1 Havcr2, Vsir* and *Tnfrsf9* in tumor infiltrating T, NKT and gd T cells of KPC3 (MRTX1133 experiments) and KPC2 (RMC6236 experiments) PDAC. **(K)** Schematic representation of orthotopic injection and MRTX1133, aLag3, aTim3, aVista and a41-BB agonist antibody treatments of KPC2 mice. **(L)** Kaplan-Meier survival curves of Veh/Iso (n=17), MRTX1133 (n=7), MRTX1133+aLag3 (n=5), MRTX1133+aTim3 (n=5), MRTX1133+aVista (n=5), MRTX1133+a41-BB agonist (n=5) treated KPC2 mice. Note: The Veh/Iso and MRTX1133 treated mice are repeated from Fig. 1H. Data are presented as mean in **B** and **D**, and as violin plots in **G, H, I** and **J**. Significance was determined by log-rank test in L. \*\*\*\**P* < 0.0001; *ns*, not significant. See related figs. S5-S6.

Next, we analyzed the transcriptional profiles of the CD8^+^ T cell infiltrates in PDACs of mice treated with Kras* targeting in combination with anti-CTLA4 therapy. Analysis of the DRGs identified downregulation of *H3f3b, Nr4a3* and *Nr4a2* in the CD8^+^ T cells. *H3f3b* has been shown to promote deletional tolerance of CD8^+^ T cells ^64^. *Nr4a3* has been shown to limit the generation of memory response^65^ and *Nr4a2* limits stem-like phenotype of the CD8^+^ T cells^66^, all indicating that the combination therapy reversed tolerance and promoted memory and stem-like CD8s. In addition, we found upregulation of 4-1BB (*Tnfrsf9*) following MRTX1133 treatment and suppression following addition of anti-CTLA4 therapy (**Supplementary Fig. S6C**), suggesting that it could be a potential target for immunotherapy. Consistent with the effector CD8 phenotype, we identified upregulation of markers of activated state including *Sp100*, *Prkcq*, Cyclin D3 (*Ccnd3*)^67–69^ and interferon responsive genes *Mndal*, *Ifi209*, *Ifi206*, *Ifi203* and *Ifi2712a* ^70^(**Supplementary Fig. S6C**) in the MRTX1133 and anti-CTLA4 treated CD8^+^ T cells. Analysis of the CD8^+^ T cells infiltrates with the RMC-6236 treated tumors further validated suppression of *H3f3b, Nr4a3, Nr4a2* and upregulation of interferon responsive genes in combination with anti-CTLA4 treatment (**Supplementary Fig. S6D**). Overall, Kras* targeting in combination with anti-CTLA4 decreased negative regulators of memory and stem-like CD8^+^ T cells and promoted interferon responsive genes indicating an activated CD8^+^ T cell state.

Next, we probed the reasons for lack of response to anti-PD1 therapy in combination with Kras* targeting despite the recruitment of exhausted CD8^+^ T cells. Analysis of T cell populations from the MRTX1133 and RMC6236 experiments indicated high levels of expression of multiple immune checkpoint genes including *Ctla4*, *Lag3*, Tim-3 (*Havcr2*), Vista (*Vsir*) and *Tnfsf9* (4-1BB) in addition to PD1 (*Pdcd1*) in the exhausted CD8^+^ T cells (**Fig. 3J**). We hypothesized that targeting the other immune checkpoints may reverse the exhaustion of the tumor infiltrating CD8^+^ T cells following Kras* targeting. Therefore, we individually treated KPC2 PDAC bearing mice with anti-Lag3, anti-Tim3, anti-Vista and anti-4-1BB (agonist antibody) in combination with MRTX1133 (**Fig. 3K**). MRTX1133 prolonged the survival of the KPC2 tumor bearing mice, but the combination of the above-mentioned checkpoint immunotherapies did not result in additional survival benefit or superior tumor inhibition in KPC2 mice (**Fig. 3K-L, Supplementary Fig. S6E-G**). Failure of other checkpoint inhibitors to reverse the exhaustion of CD8^+^ T cells indicates that these cells are likely terminally exhausted and other methods of priming are required. Our results indicate that anti-CTLA4 specifically synergizes with Kras* targeting therapies with expansion of memory/stem-like and effector CD8^+^ T cells, whereas addition of anti-PD1, anti-Tim3, anti-Lag3, anti-Vista and anti-4-1BB (agonist antibody) failed to reverse CD8 exhaustion in mice with PDAC.

### Kras* targeting in combination with anti-CTLA4 drives epigenetic reprogramming of Tregs

We next analyze the chromatin regulatory elements of sorted CD45^+^ immune cells from KPC2 PDAC mice treated with MRTX1133 in combination with anti-CTLA4 or anti-PD1 (**Fig. 4A**) by single cell ATAC sequencing (scATAC seq). scATAC seq of tumor infiltrating CD45^+^ cells identified 9 immune cell clusters including CD4^+^ T cells, Treg, CD8^+^ T cells, DCs, 3 myeloid clusters including two macrophage populations, and B cells in PDAC (**Fig. 4A**). Further analysis of the T and NK cells identified 9 T and NK cell subclusters including NK cells, CD8^+^ effector T cells, memory/stem-like CD8^+^ T cells, CD8^+^ exhausted T cells, CD8^+^ tissue-resident memory T cells (CD8 TRM), T regs, Th2 cells, naïve CD8^+^ and CD4^+^ T cells (**Fig. 4A, Supplementary Fig. S7A**). Consistent with our results from scRNA seq. experiments, Kras* targeting increased T cell infiltrates and the anti-CTLA4 combination further increased frequencies of CD8 effectors, memory/stem-like CD8^+^ T cells, naïve CD8^+^ and CD4^+^ T cells with a decrease in CD8^+^ exhausted T cell frequencies (**Fig. 4A, Supplementary Fig. S7B**). The scATAC seq method enabled identification of transcription factor (TF) activity and their fate/cell-specific binding programs in the Tregs. The suppression of cytokines IL10, IL35 and TGF-β1 is critical for limiting CD8 cytotoxicity in the tumor by Tregs, as observed in mice treated with Kras* inhibitor and anti-CTLA4 (**Fig. 3E-I**). We further evaluated TF motifs that specifically bind to the promoter/ enhancer regions to regulate the effector Treg response (**Fig. 4B, Supplementary Fig. S7C**). Analysis of the IL10, IL35 and Tgf-β genes revealed significant suppression of the peaks in the Ebi3 (IL35 component) gene promoter region and in the enhancer region of Il10 (**Fig. 4B, Supplementary Fig. S7C**). Next, we probed for TFs that can bind to the motifs in the promoter and enhancer regions of IL10 and IL-35 of Tregs. Comparison of motif enrichment of the TFs between different treatment groups in the Tregs revealed significant downregulation of Bach2, Nfatc1, Nr4a2, Maf and AP-1 related TFs, Fos, Fosb, Junb, Jund, in the Kras* inhibitor and anti-CTLA4 arm (**Fig. 4C**). The AP-1 TFs specifically bind to the promoter region of IL-35 gene to promote effector Treg function^71^. Footprint analysis of Fos, Fob, Junb and Jund binding chromatin regions revealed preferential binding of the chromatin regions in Tregs all groups compared to the MRTX1133 and anti-CTLA4 treated mice (**Fig. 4D**). Motif enrichment and footprint analysis in the Tregs revealed downregulation of the TFs at the promoter/enhancer regions of IL10 and IL35 genes involved in effector Treg differentiation in mice treated with MRTX1133 and anti-CTLA4 immunotherapy in PDAC.

**Figure 4:**
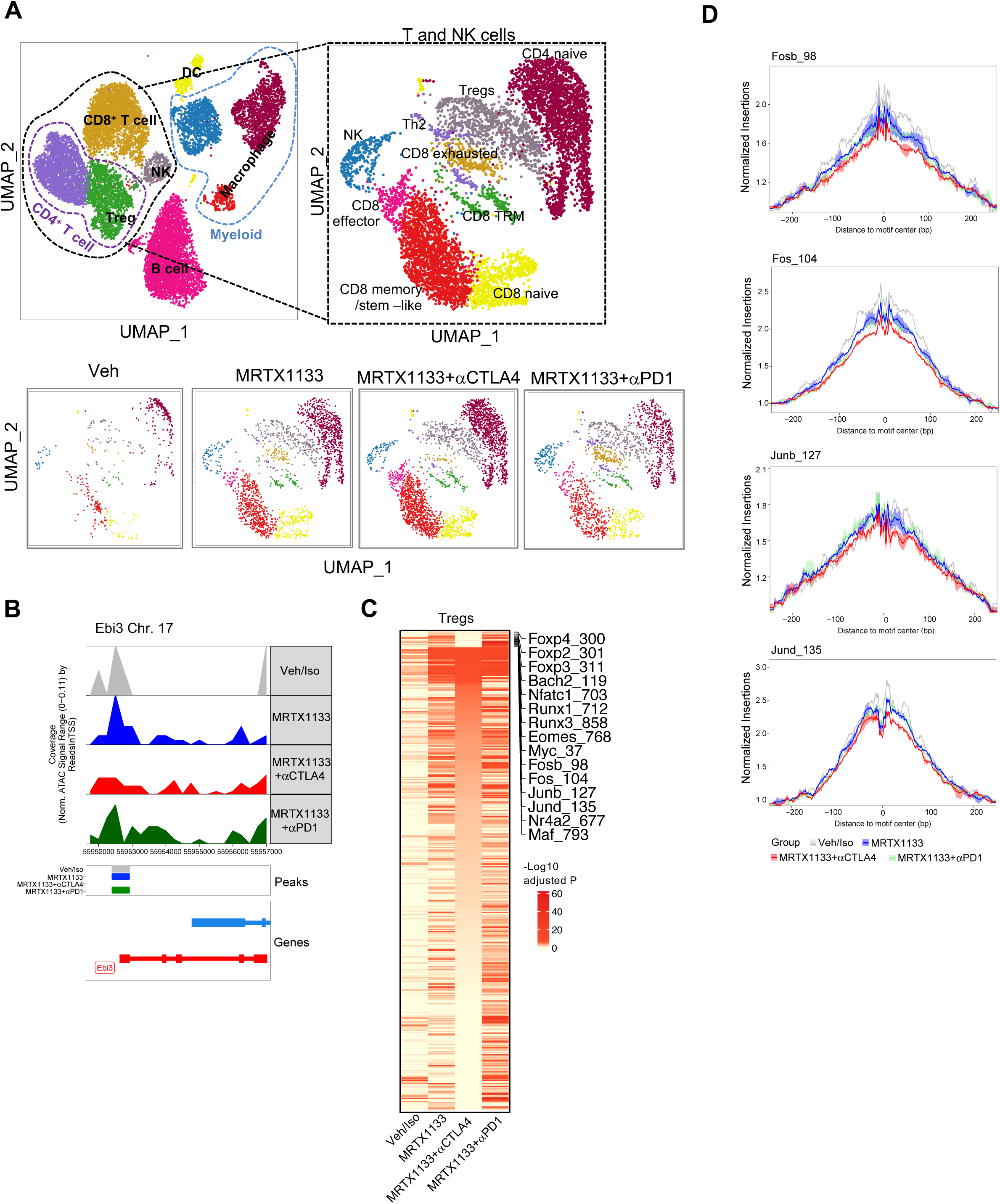
scATAC seq. analysis of tumor infiltrating Tregs in MRTX1133 and checkpoint immunotherapy treated PDAC. **(A)** UMAP of all CD45^+^ cell types (top left) and T cell subsets in Veh/Iso, MRTX1133, MRTX1133+αCTLA4, MRTX1133+αPD1 (top right, bottom) treated KPC2 PDAC (n=3 tumors/ group). **(B)** Genome browser tracks of Tregs around the Ebi3 gene locus in Veh/Iso, MRTX1133, MRTX1133+αCTLA4, MRTX1133+αPD1 treated KPC2 PDAC. **(C)** Heatmap representation of ATAC-seq motif enrichment of TFs across Tregs of Veh/Iso, MRTX1133, MRTX1133+αCTLA4, MRTX1133+αPD1 treated KPC2 PDAC. **(D)** ATAC seq. footprint aggregate plot around the predicted Fos, Fosb, Jund, and Junb motifs in Tregs of Veh/Iso, MRTX1133, MRTX1133+αCTLA4, MRTX1133+αPD1 treated KPC2 PDAC. See related figs. S7.

### Depletion of CD4^+^ T cells or CD25^+^ Tregs synergize with Kras* targeting in PDAC

The transcriptional changes observed in tumor-infiltrating Tregs, CD4⁺, and CD8⁺ T cells from mice treated with Kras* targeting combined with anti-CTLA-4 therapy prompted us to investigate the functional roles of these T cell subsets in the immune regulation of PDAC (**Fig. 3, 4**). Anti-CTLA4 combination with MRTX1133 and RMC6236 did not result in a significant Th1 response (**Fig. 3A-D**) (as marked by expansion of CD4^+^ Tbet^+^ cells in response to αCTLA4 therapies^72^) despite robust PDAC inhibition and effector CD8 response. To determine the functional contribution of the T regs, CD4^+^ T cells, and to assess the contribution of ‘CD4 help’ in generating effector CD8 responses, we treated the KPC2 tumor bearing mice with anti-CD25 and anti-CD4 depletion antibodies in combination with MRTX1133 therapy (**Fig. 5A**). Analysis of peripheral blood confirmed systemic depletion of CD4^+^ T cells and CD25^+^ T cells in the anti-CD4 and anti-CD25 antibodies treated mice (**Supplementary** Fig. 8A-B). Although depletion of CD4^+^ T cells did not result in significant changes in other T cell populations, depletion of CD25^+^ T cells resulted in an increased frequencies of CD4^+^ CD25^−^ Foxp3^+^ population in the circulation (**Supplementary** Fig. 8A-B). MRTX1133 treated KPC2 mice succumbed to PDAC despite prolonged survival (**Fig. 5B**). Depletion of CD25^+^ cells in combination with MRTX1133 further improved the survival of the KPC2 mice (**Fig. 5B**). Complete depletion of the CD4^+^ T cells resulted in robust inhibition of PDAC in combination with MRTX1133 with similar survival kinetics as the MRTX1133 + anti-CTLA4 treated mice (**Fig. 5B**).

**Figure 5:**
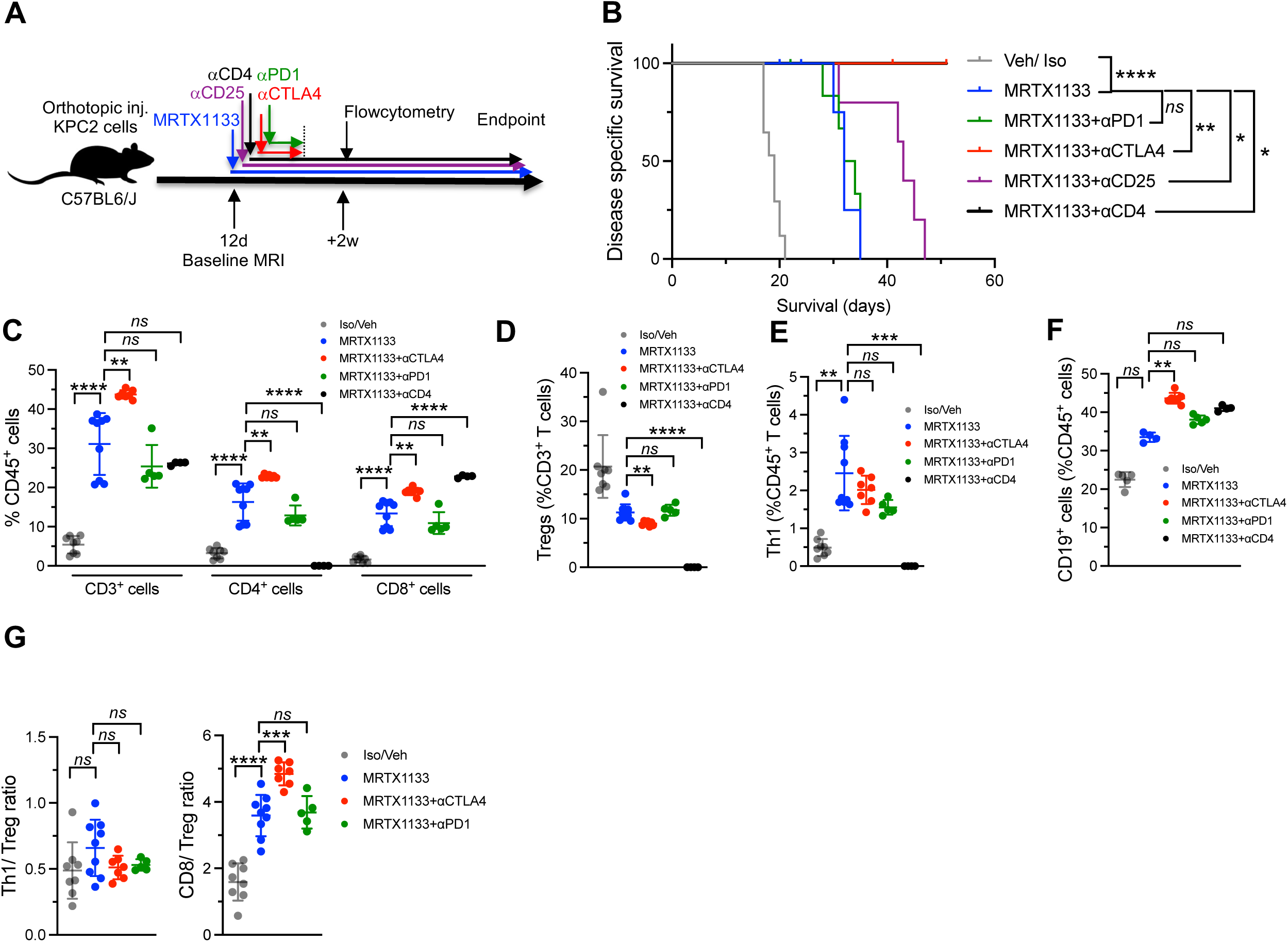
Depletion of CD4^+^ T cells or Tregs synergize with Kras* targeting in PDAC. **(A)** Schematic representation of orthotopic injection and MRTX1133, aCD4 and aCD25 treatments of KPC2 mice. **(B)** Kaplan-Meier survival curves of Veh/Iso (n=17), MRTX1133 (n=11), MRTX1133+aCD4 (n=5), MRTX1133+aCD25 (n=10), MRTX1133+ aCTLA4 (n=11), MRTX1133+aPD1 (n=9) treated KPC2 mice. <u>Note:</u> The Iso/Veh, MRTX1133, MRTX1133+ aCTLA4, and MRTX1133+aPD1 curves are repeated from Fig. 2B to allow for direct comparisons. **(C-G)** Immunophenotyping analysis of KPC2 PDAC of indicated groups. CD3^+^(T cells); CD3^+^CD4^+^ (CD4^+^ T cells); CD3^+^CD8^+^ (CD8^+^ T cells) were measured as a percentage of CD45^+^ cells **(C)**; CD4^+^Foxp3^+^ Tregs were measured as a percentage of CD3^+^ cells **(D)**; CD4^+^T-bet^+^ (Th1) measured as a percentage of CD45^+^ cells **(E)**, and CD19^+^ B cells measured as a percentage of CD45^+^ cells **(F)**. CD8/Treg ratio and Th1/Treg ratio **(G)**. n=4-9 tumors/ group. Data are presented as mean ± SD in **C, D, E, F** and **G**. Significance was determined by log-rank test in **B**, Kruskal-Wallis with Dunn’s multiple comparisons test in **E**, one-way ANOVA with Dunnett’s multiple comparisons test in **C, D, F** and **G**. \**P* < 0.05, \*\**P*< 0.01, \*\*\**P* < 0.001, \*\*\*\**P* < 0.0001; *ns*, not significant. See related figs. S8-S9.

Next, to evaluate mechanisms underpinning the alterations in tumor infiltrating T cells as a result of immune checkpoint blockade therapies/T cell depletions, we performed flowcytometry analysis in the age-matched KPC2 tumors treated with MRTX1133 in combination with anti-PD1, anti-CTLA4 and anti-CD4 depletion antibodies. Consistent with the findings from scRNA seq. analysis, MRTX1133 treated resulted in an increase in CD3^+^ T, CD3^+^ CD4^+^ T, CD3^+^ CD8^+^ T and CD4^+^ Foxp3^+^ T cell frequencies in the PDAC TME (**Fig. 5C**). The addition of anti-CTLA4 antibody to MRTX1133 treated PKC tumors demonstrated further increase in total T cells, CD4^+^ T cells and CD8^+^ T cell frequencies (**Fig. 5C**). The anti-CD4 antibody treated mice in combination with MRTX1133 demonstrated depletion of intra-tumoral CD4^+^ T cells, T regs and enhanced CD8^+^ T cell frequencies further validating that the CD4^+^ T cells are inhibitory to the CD8^+^ T cell function in PDAC (**Fig. 5C, D**). Analysis of the Th1 response in PDAC revealed that the Th1 cells were a rare population in the TILs (∼1%), which increased following Kras* targeting with MRTX1133 (∼2%). However, no further changes in the Th1 population was observed following addition of anti-CTLA4 therapy (**Fig. 5E**). Further analysis of the B cell frequencies demonstrated a significant increase in B cell frequencies in mice that received MRTX1133 and anti-CTLA4 therapy (**Fig. 5F**). In addition, analysis of Th1/Treg ratios did not demonstrate any differences between the treatment groups (**Fig. 5G**). Analysis of the CD8/Treg ratio demonstrated an increase with MRTX1133 therapy and further increase with addition of anti-CTLA4 therapy indicating a suppression of T regs with enrichment of CD8 response (**Fig. 5G**). Collectively, robust tumor inhibition response and flowcytometry analysis following CD4^+^ T cell depletion indicates that CD4^+^ T cells are collectively inhibitory to CD8^+^ T cells and that the ‘CD4 help’ is dispensable in the immune regulation of PDAC.

To explore whether our findings are unique to PDAC and to confirm that the mechanisms of T cell regulation in combination with anti-CTLA4 and anti-PD1 therapy are distinct in PDAC infiltrating lymphocytes, we repeated the earlier experiments used to identify the mechanism of action of anti-CTLA4 and anti-PD1 immunotherapies using the checkpoint responsive MC-38 colorectal cancer (CRC) model ^72^. We treated the MC-38 tumor bearing mice with anti-CTLA4 and anti-PD1 checkpoint immunotherapies and performed flowcytometry analysis at similar time points described in earlier studies (**Supplementary Fig. S9A**)^72^. Consistent with earlier studies, both anti-CTLA4 and anti-PD1 antibodies suppressed tumor growth (**Supplementary Fig. S9B-D**), albeit through different immune regulatory mechanisms (**Supplementary Fig. S9E-H**). The anti-CTLA4 therapy resulted in an increase in frequencies of CD4^+^ Tbet^+^ (Th1) cells, CD8^+^ T cells, CD8^+^ Tbet^+^ effector T cells, higher Th1/Treg and CD8/T reg ratios, whereas the αPD1 therapy resulted in predominantly a CD8^+^ T cell response (**Supplementary Fig. S9E-H**). These control experiments further strengthen our conclusion that anti-CTLA4 therapy synergizes with Kras* targeting through inhibition of T regs and reversing exhaustion of CD8^+^ T cells in PDAC, a mechanism that is both distinct from inhibition of other immune checkpoints in PDAC and is also unique to this tumor type. In contrast to antiCTLA-4 blockade in CRC, there was no appreciable Th1 response with the addition of αCTLA4 to Kras* targeting therapies and the CD4 help is dispensable for effector CD8 responses in PDAC.

### Kras* targeting in combination with **α**CTLA4 induces formation of tertiary lymphoid structures

In depth histological analysis of PDAC tissues from KPC2 mice demonstrated an increase in the presence of tertiary lymphoid structures (TLS) in mice treated with MRTX1133 or RMC6236 in combination with anti-CTLA4 (**Fig. 6A-B, Supplementary Fig. S10A, and S11A-B**). Although occasional TLS were found in PDAC tissues of mice treated with MRTX1133/ RMC6236 monotherapy or in combination with αPD1 antibody, the combination of Kras* targeting with anti-CTLA4 demonstrated consistent increase TLS with 100% penetrance in the KPC2 tumors (**Fig. 6A-B, Supplementary Fig. S10A, S11A-B**). Further, histological assessment of age-matched KPC and KPPC GEMs treated with MRTX1133 and anti-CTLA4 therapy revealed TLS with 100% penetrance in mice that received combination therapy (**Fig. 6C-F, Supplementary Fig. S12A-B**). TLS refers to the *de novo* occurrence of organized aggregates of immune cells in non-lymphoid organs^73,74^. In the context of cancer and immunotherapy, the presence of TLS is frequently associated with favorable responses to immunotherapy and better prognosis ^73,74^. The composition of TLS may vary depending on the host tissue but are frequently characterized by an inner zone of B cells, surrounded by T cells and accompanied by the presence of follicular dendritic cells (FDCs) which enable formation of germinal center B cells, follicular T helper (T_FH_) cells and mature dendritic cells^73–77^. In addition, the TLS are accompanied by endothelial venules that provide the associated vasculature that likely mediate lymphocyte recruitment ^77^. To confirm the histopathological identification of TLS, we performed immunostaining analysis for markers of T cells, B cells, endothelial vasculature, FDCs and T_FH_ cells (**Fig. 6G-L**). Immunofluorescence staining with Tyramide signal amplification (TSA) multiplex analysis revealed presence of CD19^+^ B cells, CD8^+^ T cells, CD31^+^ endothelial cells in vascular structures and CD21^+^ and CD23^+^ cells (expressed by T_FH_, FDCs and B cell populations) confirming that these immune aggregates seen with Kras* targeting and αCTLA4 combination therapy are TLS at varying stages of maturation (**Fig. 6G-L**). We ascertain that each lymphoid structure that was identified as TLS demonstrated neo-vascularization with formation of endothelium identified by CD31 staining with co-existence of T and B cell regions in orthotopic KPC2 mice and KPC, KPPC GEMs that received combination therapy (**Supplementary Fig. S13A-B, S14A-B**).

**Figure 6:**
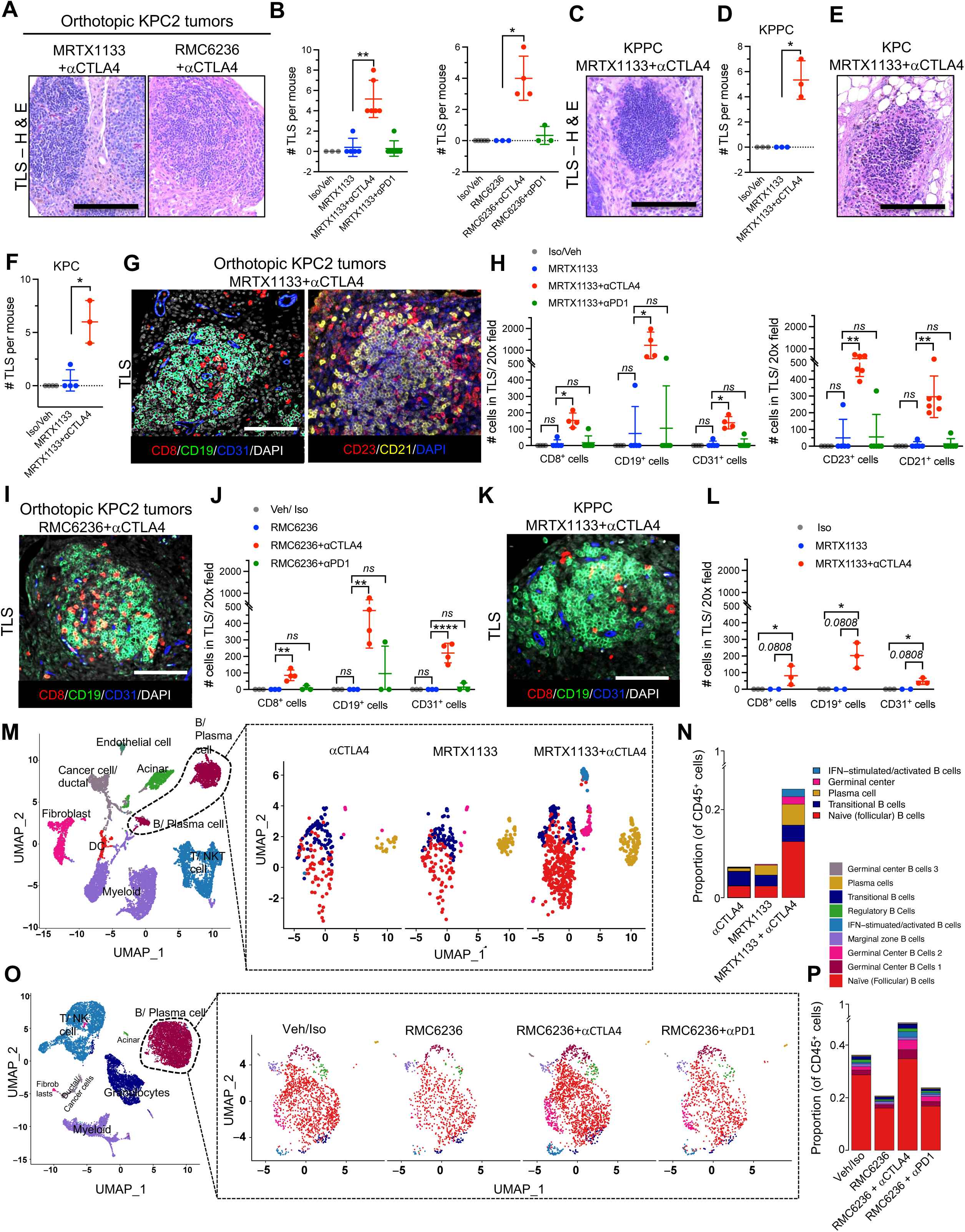
Kras* targeting in combination with αCTLA4 induces formation of tertiary lymphoid structures in PDAC. **(A-B)** Representative H&E sections of TLS in MRTX1133, RMC6236 treated KPC2 mice with aCTLA4/ aPD1 **(A)**, with quantification of number of TLS (n=3-7 tumors/ group) **(B)**. **(C-D)** Representative H&E sections of TLS in MRTX1133 treated KPPC mice with aCTLA4 **(C)**, with quantification of number of TLS (n=3 tumors/ group) **(D)**. **(E-F)** Representative H&E sections of TLS in MRTX1133 treated KPC mice with aCTLA4 **(E)**, with quantification of number of TLS (n=3-4 tumors/ group) **(F). (G-H)** Representative CD8-CD19-CD31-DAPI (left) and CD21-CD23-DAPI (right) immunofluorescence images of TLS **(G)**, with quantification of cell numbers in the TLS regions of MRTX1133 treated KPC2 mice with aCTLA4/ aPD1 **(H)** (n=4-6 tumors/ group). **(I-J)** Representative CD8-CD19-CD31-DAPI immunofluorescence images of TLS **(I)**, with quantification of cell numbers in the TLS regions of RMC6236 treated KPC2 mice with aCTLA4/ aPD1 **(J)** (n=3-4 tumors/ group). **(K-L)** Representative CD8-CD19-CD31-DAPI (left) and CD21-CD23-DAPI (right) immunofluorescence images of TLS **(K)**, with quantification of cell numbers in the TLS regions of MRTX1133 treated KPC2 mice with aCTLA4/ aPD1 **(L)** (n=2-3 tumors/ group). **(M-N)** UMAP of all cell types (left) and B cell subsets in aCTLA4, MRTX1133, MRTX1133+aCTLA4 treated (right) KPC3 PDAC **(M)**, with number of cells in B cell subsets (as % of CD45^+^ cells) **(N)** determined by scRNA seq. analysis (n=3 mice/group). **(O-P)** UMAP of all cell types (left) and B cell subsets in Veh/Iso, RMC6236, RMC6236+aCTLA4, RMC6236+aPD1 treated (right) KPC2 PDAC **(O)**, with number of cells in B cell subsets **(P)** determined by scRNA seq. analysis (n=3 tumors/group). <u>Note:</u> B cells from UMAPs are analyzed in **Fig. 6M** and **Fig. 6O** and T cells from the scRNA seq. data are analyzed in **Fig. 3A** and **3C** respectively. Data are presented as mean in **N** and **P**, mean ± SD in **B, D, F, H, J** and **L**. Significance was determined by Kruskal-Wallis with Dunn’s multiple comparisons test in **B, D, F, H** and **L**, one-way ANOVA with Dunnett’s multiple comparisons test in **J**. \**P* < 0.05, \*\**P*< 0.01, \*\*\*\**P* < 0.0001; *ns*, not significant. Scale bars-100mm. See related figs. S10-S15.

Next, we performed single cell RNA sequencing (scRNA seq.) analysis of the sorted live cells from KPC3 PDAC bearing mice treated with MRTX1133 and/ or anti-CTLA4 antibody (**Fig. 6M**). UMAP analysis of the scRNA seq. dataset captured 8 clusters including fibroblasts, cancer/ductal cells, acinar cells, myeloid cells, DCs, T/NKT cells, endothelial cells and B/plasma cells (**Fig. 6M**). Further sub-clustering of the B and plasma cell populations revealed an increase in the frequencies and number of naïve (follicular B cells) (expressing *Ighd*, *Ms4a1*, *Pax5*, *Cd21* and *Cd23*), plasma cells (expressing *Igkc*, *Xbp1*, *Jchain*, *Tnfrsf13b*, *Sdc1*, *Tnfrsf17*, *Cd320*, *Irf4* and *Prdm1*), germinal center B cells (*Aicda*, *Gesam*, *S1pr2*, *Bcl6* and *Fas*) and IFN-stimulated/ activated B cell populations (*Ifit3*, *Slfn5* and *Herc6*) (**Fig. 6M-N, Supplementary Fig. S15A-C**). To validate our findings from the MRTX1133 and anti-CTLA4 combination therapy, we used a second model with the RMC6236 compound in combination with anti-CTLA4 and anti-PD1 therapy. We performed scRNA seq. analysis on sorted live, CD45^+^ cells to enrich for the immune populations from PDAC of age-matched RMC6236 mice treated KPC2 mice in combination with anti-CTLA4 and anti-PD1 therapy (**Fig. 6O-P**). UMAP analysis of the scRNA seq. dataset captured 4 major clusters including T/NK cells, granulocytes, myeloid cells and B/plasma cells and 3 clusters with residual populations of acinar cells, fibroblasts and ductal/cancer cells (**Fig. 6O**). Further sub-clustering of the B/plasma cell populations revealed increase in naïve (follicular) B cells, germinal center B cells (3 clusters) and IFN-stimulated/ activated B cell populations in the RMC6236 and anti-CTLA4 treated tumors (**Fig. 6O-P, Supplementary Fig. S15D-F**). The plasma cell populations weren’t captured in the RMC6236 treated tumors as they are negative/ low for CD45 (**Fig. 6M-P**). Interestingly, minor populations of regulatory B cells and marginal zone B cells were observed in the CD45^+^ sorted samples that weren’t captured in the earlier analysis of B cells. However, treatment with RMC6236, anti-CTLA4/ anti-PD1 did not affect these cell numbers in the KPC2 tumors (**Fig. 6O-P**). Further analysis of the transcriptional profiles of B cells in the Kras* targeting and anti-CTLA4 therapy indicated that the IFN stimulated/ activated B cells demonstrate an increase in cytokines including hallmark interferon-stimulated genes viz. *Ifit3*, *Slfn5*, and *Isg15* and germinal center B cell transcription factor *Bcl6* (**Supplementary Fig. S15A, D**). Our data establishes the recruitment of TLS in mice treated with Kras* targeting MRTX1133/RMC6236 in combination with anti-CTLA4 in multiple orthotopic and autochthonous GEMs. The presence of neo-vascularization, specific T and B cell recruitment, expression of CD21 and CD23, expansion of follicular B cells, plasma cells and IFN-stimulated/activated B cells validate that these structures are TLS and not lymphoid aggregates.

### Tertiary lymphoid structures functionally contribute to synergy of Kras* targeting with anti-CTLA4 therapy

Next, to determine whether the TLS play a functional role in mediating synergy between MRTX1133/RMC6236 and anti-CTLA4 in PDAC, we treat mice that received combination therapy with lymphotoxin-β inhibitor (LTBi) (Baminercept) as it was the most robust cytokine expressed by the tumor infiltrating B cell (**Fig. 7A, Supplementary Fig. S16A-B**). In addition, we use an antibody to deplete CD19^+^ B cells in mice that received combination therapy (**Fig. 7A, Supplementary Fig. S16C-D**). Analysis of peripheral blood from mice that received anti-CD19 depletion antibody revealed systemic depletion of CD19^+^ B cells **(Supplementary Fig. S16C-D)**. LTBi and B cell depletion reversed the survival benefit conferred by the combination, despite marginal increase in survival compared to mice that received MRTX1133/ RMC6236 monotherapy indicating that the TLS are functionally critical for synergy of Kras* inhibitors with anti-CTLA4 therapy (**Fig. 7B-F**). We performed histological analysis of the pancreatic tissues when the mice that received MRTX1133/ RMC6236 monotherapy, MRTX1133/ RMC6236 in combination with anti-CTLA4 and LTBi, and MRTX1133/ RMC6236 in combination with anti-CTLA4 and anti-CD19 succumbed to PDAC (**Fig. 7D-F**). The mice that received MRTX1133/ RMC6236 and anti-CTLA4 demonstrated robust tumor inhibition at a time point when the tumors with inhibition of TLS succumbed to PDAC (**Fig. 7D-F**). Our data indicates the functional contribution of TLS in PDAC inhibition with combination therapy (**Fig. 7D-F**). Our data establishes that Kras* targeting resulted in recruitment of T cell infiltrates and synergizes with anti-CTLA4 therapy but not with other checkpoint inhibitors including anti-PD1, anti-Tim3, anti-Lag3, anti-Vista or anti-4-1BB agonist antibody therapy. The combination of Kras* targeting and anti-CTLA4 therapy resulted in recruitment of TLS in the pancreatic TME with expansion of naïve (follicular) B cells, plasma cells, germinal center B cells and IFN-stimulated/ activated B cells (**Fig. 8A**).

**Figure 7:**
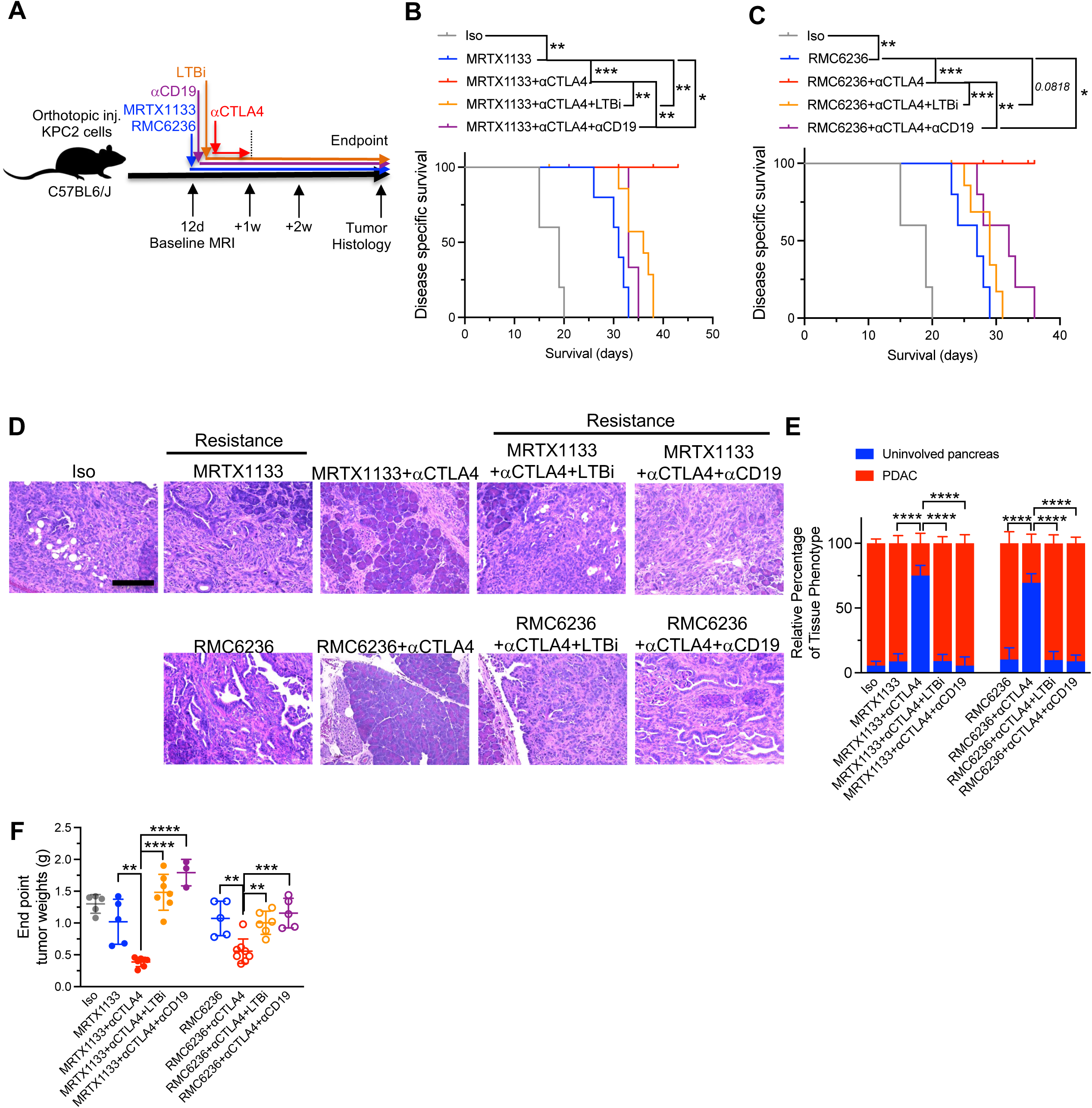
Inhibition of tertiary lymphoid structures negates the effects of Kras* targeting and anti-CTLA4 combination therapy. **(A)** Schematic representation of orthotopic injection and MRTX1133, RMC6236, aCD19, aCTLA4 and LTBi (Baminercept) treatments of KPC2 mice. **(B-C)** Kaplan-Meier survival curves of Iso (n=5), MRTX1133 (n=5), MRTX1133+αCTLA4 (n=8), MRTX1133+aCTLA4+aCD19 (n=4), MRTX1133+αCTLA4+LTBi (n=8) **(B)**, and RMC6236 (n=5), RMC6236+αCTLA4 (n=8), RMC6236+aCTLA4+aCD19 (n=5), RMC6236+αCTLA4+LTBi (n=8) **(C)** treated KPC2 mice. <u>Note:</u> Same Iso treated mice were repeated in Fig. 7B and 7C to allow for direct comparisons. **(D-E)** Representative H&E sections in MRTX1133, RMC6236, aCD19, aCTLA4 and LTBi treated KPC2 mice **(D)**, with quantification of relative percentage of tissue phenotype (n=3-7 tumors/ group) **(E). (F)** Endpoint tumor weights of mice from the indicated treatment groups (n= 3-8 mice/group). Data are presented as mean ± SD in **E** and **F**. Significance was determined by two-way ANOVA in **E**, log-rank test in **B** and **C**, and by one-way ANOVA with Sidak’s multiple comparisons test in **F**. \**P* < 0.05, \*\**P*< 0.01, \*\*\*\**P* < 0.0001. Scale bars-100μm. See related fig. S16.

**Figure 8:**
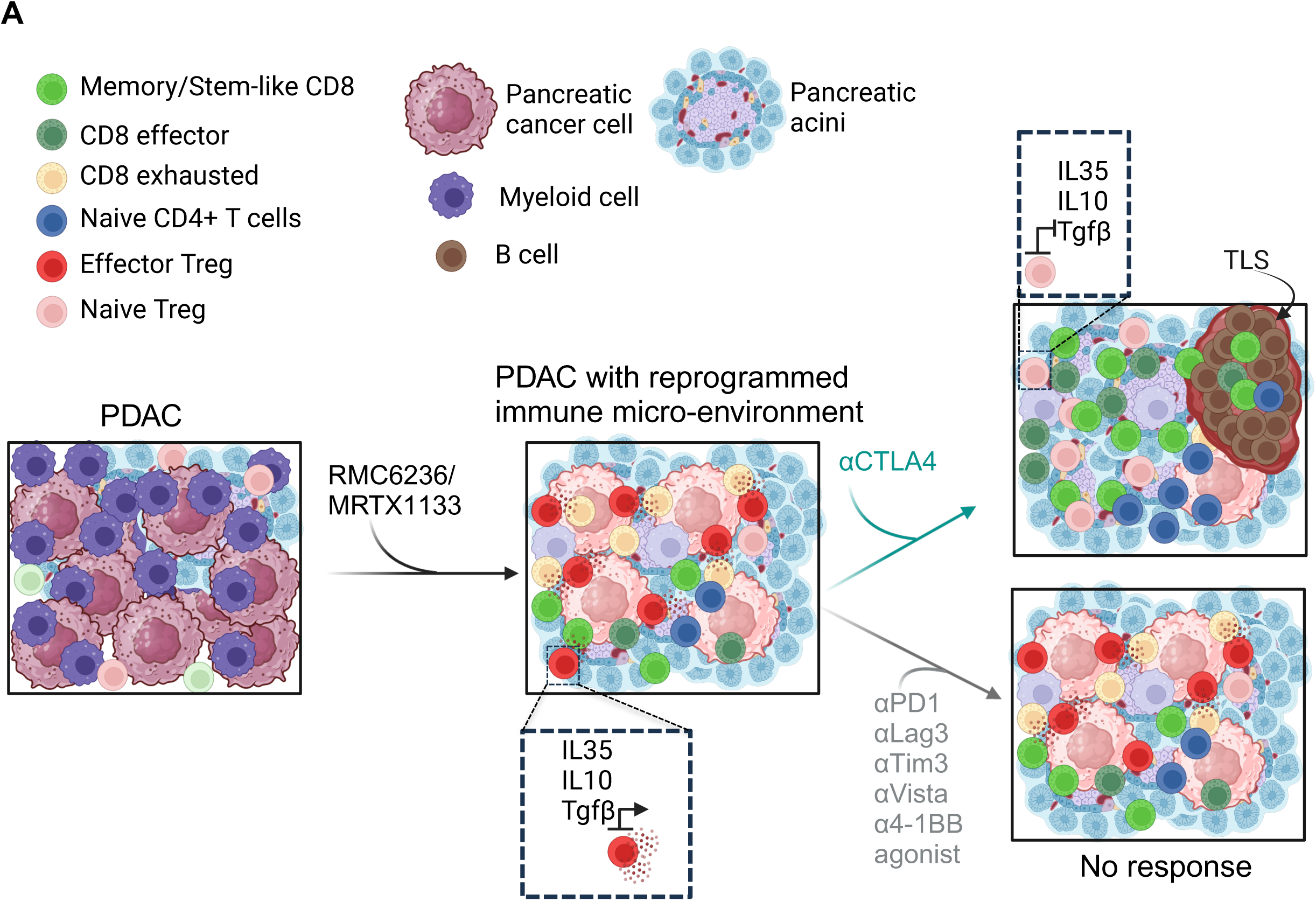
(A) Graphical abstract summarizing the synergy between Kras* targeting and checkpoint immunotherapy.

## Discussion

Pancreatic cancer remains refractory to checkpoint immunotherapies in pre-clinical models and clinical studies ^45,46,78,79^. Genetic silencing and pharmacological inhibition of Kras* results in an increase in tumor infiltrating CD4^+^ T cells, CD8^+^ T cells, Tregs and B cells in the acute phase with complete regression of advanced PDAC in genetic models^40,41^. However, influx of immune cells did not result in a long-term memory/ activated CD8^+^ T cell response and therefore led to escape from pharmacological Kras* targeting with mice succumbing to PDAC. Our results demonstrate that irrespective of the Kras* targeting strategy, specific addition of anti-CTLA4 combination led to long-term PDAC suppression, inhibition of effector Tregs, switch to a naïve Treg phenotype, enhanced CD8 memory and effector response in combination with development of TLS within the PDAC. The suppression of the effector Treg cytokines Il10, Tgfb1, and Il35 contributed to the naïve Treg phenotype and expansion of the effector CD8 response in the tumors that received anti-CTLA4 combination therapy. Our data indicates that anti-CTLA4 combination epigenetically suppresses II35 and Il10 in Tregs likely through downregulation of AP-1 family of transcription factors Fos, Fosb, Junb, and Jund at the promoter and enhancer regions of these genes.

It is interesting to note that targeting Kras* results in reprogramming of the tumor microenvironment reflecting all its constituents including the immune cells^80^. This observation highlights that Kras* mediated oncogenesis significantly reprograms the microenvironment towards tumor immune responses with causal impact. Deletion of CD4^+^ T cells or CD25^+^ T cells enhances the impact of Kras* inhibitors, directly implicating the need for productive immune responses in realizing the full efficacy of Kras* inhibitors. Targeting Kras* results in cancer cells alters many signaling pathway axis and likely blocks secretome that inhibits recruitment of T cells and B cells. Consequently, Kras* targeting favors secretome that can help their recruitment with functional impact. Our data suggest that while the resulting immune responses is of benefit, it is not optimal and therefore leads to PDAC relapse. But if relevant immune checkpoint inhibitor is combined with Kras* targeting, long-term benefits are observed without emergence of in majority of the mice.

Analysis of exhausted CD8^+^ T cells following Kras* targeting revealed high expression of multiple immune checkpoint genes including Tim3, Lag3, 4-1BB, PD1 and Vista. Interestingly, treatment with individual checkpoint therapies such as anti-Tim3, anti-Lag3, anti-41-BB agonist antibody, anti-Vista and anti-PD1 did not reverse CD8^+^ T cell exhaustion in combination with the Kras* inhibitors. A likely explanation could be that these CD8^+^ T cells are terminally exhausted and unable to reverse with individual checkpoint targeting (*vide supra*), and additional methods of priming are required to recruit new CD8^+^ T cells^53^. Our results indicate that addition of CTLA4 blockade likely results in an expansion of tumor reactive CD8^+^ T cells and broadens the T cell repertoire in addition to reversing existing exhausted CD8 populations^81–83^. Surprisingly, another study using subcutaneous PDAC model demonstrates marginal improvement in survival with Kras* targeting in combination with anti-PD1 therapy^84^. However, this study represents a PDAC setting with intrinsically high T cell infiltration and sensitivity to checkpoint blockade as a monotherapy^50–52,84^. Further, subcutaneous tumor models lack the tumor microenvironment and associated lymphoid tissue unique to organs in the gastrointestinal tract. Our work determines the role of immune checkpoint inhibitors in a cancer context with the appropriate PDAC tumor microenvironment, lymphatic drainage and stromal components in pancreatic cancer. Another study using a genetically engineered iKPC PDAC model with long-term follow up demonstrated that Kras^G12D^ inhibition does not synergize with anti-PD1 therapy^85^, consistent with our findings indicating that combination therapy with anti-PD1 did not reverse terminally exhausted CD8s in PDAC.

Further analysis of the Tregs/ CD4^+^ T cells with depletion of specific cell populations revealed that the CD4s are inhibitory overall and impede the anti-tumor CD8 response. Surprisingly, our data with multiple PDAC models demonstrate a lack of Th1 response with Kras* targeting monotherapy or with anti-CTLA4 combination therapy indicating that ‘CD4 help’ could be dispensable for the generation of effector/ memory CD8^+^ T cell response in PDAC. Comparative analysis of melanoma and PDAC infiltrates from patient samples further confirms that Th1 cells were nearly non-existent in the PDAC TME^86^. In the context of melanoma, Th1 and Tregs comprised the majority of the CD4^+^ infiltrates. Anti-CTLA4 therapy resulted in an expansion of the Th1 helper response accompanied by Treg suppression. Our prior work in this area which probed the role of Kras^G12D^ targeting on the immune microenvironment using the inducible KPC GEM models indicate that the tumor infiltrating CD4^+^ T cells differentiate into Th2, Th17 and Treg lineages and not the Th1 lineage in PDAC^40^. Depletion of CD4^+^ T cells in the context of Kras^G12D^ targeting heightened the CD8^+^ cytotoxic lymphocyte frequencies further indicating that CD4s overall are inhibitory to CD8^+^ T cell response ^40^. Other studies that probe the functional role of CD4^+^ T cells, in particular Th2 and Th17 in different PDAC contexts with pre-clinical depletion models conclude that the CD4^+^ T cells are inhibitory to the cytotoxic CD8^+^ T cell response^87–89^. Taken together, our work highlights the unique profile of the tumor infiltrating lymphocytes and the distinct anti-tumor immune responses in PDAC.

Combination therapy (Kras* targeting and CTLA4 blockade) leads to emergence of TLS in PDAC. The emergence of TLS as a response to immunotherapies represents a favorable prognostic indicator for control of cancer^73,74^. TLS occurring in melanoma patients was first identified as a marker of good prognosis and response to immunotherapies. Subsequently, the presence of TLS across multiple cancer models have been associated with longer survival rates with prolonged tumor inhibition ^90–92^.

TLS differ from the secondary lymphoid organs such as lymph nodes as they lack encapsulation, which enables direct access of the cellular components (CD8^+^ T cell, B cell and plasma cells) to facilitate PDAC inhibition^73,74,93^. It is unclear what the lymphoid tissue stimulatory components are in inducing TLS in cancer. Earlier studies have implicated several immune populations including Th17 cells^94^, some subsets of innate lymphoid cells (ILCs)^91,95^, effector CD8^+^ T cells and B cells ^96–98^. In the PDAC models here, Kras* inhibition resulted in an increase in CD4^+^ T cell infiltrates including CD4^−^ Foxp3^−^ T cells and Th17 cells as shown in prior studies^40,87^, CD8^+^ T cells (including exhausted and effector CD8s) and B cell infiltrates. However, the presence of TLS in the context of Kras* targeting occurs only in combination with anti-CTLA4 which likely indicates that an enhanced CD8 effector response with suppression of exhausted CD8s and effector T regs drives the formation of TLS.

TLS are frequently seen in cancer at different stages of maturation including immature (consisting of dense lymphocyte aggregates without markers of FDCs and B/T cell zones), primary follicle like TLS (presence of FDCs but lacking germinal centers) and secondary follicle like TLS (mature with active germinal centers) ^73,99,100^. The combination of Kras* targeting and anti-CTLA4 resulted in expansion of germinal center B cells and plasma cells with expression of FDC/ T_FH_ markers CD21 and CD23 likely indicating that the TLS were maturing with functional capacity. Another critical aspect of anti-tumor immunity associated with TLS is whether they can re-educate the T cells to accentuate an effector CD8 response. Although an increase in IFN stimulated/activated B cells and effector CD8^+^ T cells was observed with the Kras* targeting combination with anti-CTLA4 therapy, lack of expansion of the T_FH_ cells or Th17 cells in the scRNA seq. analysis of combination therapy tumors indicates that the TLS associated with PDAC aren’t fully mature, as also seen in our immunostaining analysis. To determine whether the TLS contribute functionally to PDAC inhibition with the combination therapy, we used them in combination with LTBi and B cell depletion experiments. Our results further validate that although the TLS are at varying stages of maturation, they are critical in realizing the full potential of Kras* targeting in combination with anti-CTLA4 therapy.

Overall, this study provides a detailed assessment of the adaptive immune response and T cell regulatory networks upon Kras* inhibition and provides compelling evidence for combination with ipilimumab in the clinic for long-term benefit.

## Material and Methods

### Mouse studies

Several primary PDAC cells lines, 343P KPC (*Pdx1-cre, LSL-Kras^G12D/+^, Trp53^R172H/+^; Rosa-LSL-EYFP* cell line, referred to as KPC1), Hy19636-PKC (*P48-Cre, LSL-Kras^G12D/+^, Trp53^F/+^*cell line, referred to as KPC2), KPC-T (*Pdx1cre, LSL-Kras^G12D/+^, Trp53^F/+^*, referred to as KPC3) were used for orthotopic implantation into the tail of the pancreas of 8-16w old C57BL6/J mice (concentration 0.5 x 10^6^ cells in 20μL PBS). The autochthonous KPC (*Pdx1-cre, LSL-Kras^G12D/+^, Trp53^R172H/+^*) and KPPC ((*P48-Cre, LSL-Kras^G12D/+^, Trp53^F/F^*) mice have been characterized earlier^101,102^. For experiments with immune checkpoint inhibitors, anti-mouse CTLA4 (BioXcell, clone 9H10, catalog no. BE0131) or anti-mouse PD-1 (BioXcell, clone 29F.1A12, catalog no. BE0273) were injected three times in 1 week, intra-peritoneally in a final volume of 100 μL as indicated. A starting dose of 200 μg was followed by two doses of 100 μg each (1 mg/mL concentration). Isotype treatments in this group received a combination of Rat IgG2a (BioXcell, clone 2A3, BE0089) and Syrian hamster IgG (BioXcell, BE0087) in the same route, time, and dosing as the checkpoint inhibitor antibodies. For anti-mouse Lag3 (BioXcell, clone C9B7W, catalog no. BE0174), anti-mouse Tim-3 (CD366) (BioXcell, clone, catalog no. BE0115), anti-mouse Vista (BioXcell, clone RMT3-23, catalog no. BE0310), anti-mouse 4-1BB (CD137) (BioXcell, clone LOB12.3, catalog no. BE0169) were injected intraperitoneally, 200 μg starting dose, 100 μg (2 doses) in a final volume of 100 μL PBS in a span of 1 week. The control mice received a cocktail of corresponding isotype antibodies include Rat IgG1 (BioXcell, Catalog no. BE0088), Rat IgGa2 (BioXcell, clone 2A3, catalog no. BE0089), Armenian hamster IgG (BioXcell, catalog no. BE0091) in the same route, time, and dosing as the checkpoint inhibitor antibodies. For CD4^+^ and CD25^+^ T cell depletion experiments, anti-mouse CD4 (BioXcell, clone GK1.5, catalog no. BE0003-1) or anti-mouse CD25 (IL-2Rα) (BioXcell, clone PC-61.5.3, catalog no. BE0012) and their isotype antibodies (Rat IgG2b, BioXCell, cloneLTF2, catalog no. BE0090 and Rat IgG1, BioXcell, clone HRPN, Catalog no. BE0088) were injected intraperitoneally, twice per week, 200μg of antibody for the course of the experiments in a final volume of 100μL PBS. For CD19^+^ B cell depletion experiments, anti-mouse CD19 (BioXcell, clone1D3, catalog no. BE0150) or the isotype Rat IgG2a (BioXcell, clone 2A3, BE0089) were injected intraperitoneally, 200μg of antibody, twice per week as indicated. For lymphotoxin-β inhibiton, 100μg of baminercept was injected intraperitoneally, every 5 days for the entire course of the experiments. For MC38 experiments, 0.5 x 10^6^ cells were injected in a final volume of 50 μL PBS subcutaneously in 8-10w old C57BL6/J mice. For MRTX1133 experiments, mice were treated with 30mg/kg MRTX1133 in 100 μL of 10% captisol (SelleckChem, Catalog No. S4592) in 50mM sodium citrate (pH 5) (Tekanova) vehicle BID i.p. or the corresponding vehicle in control mice as described earlier^1,41^. For RMC-6236 experiments, mice were treated with RMC-6236, prepared using formulation of 10% DMSO (MP Biomedicals, 0219605590), 20% PEG-400 (Possible Missions Inc, NC0572735), 10% Solutol HS15 (SelleckChem, Catalog No. E1708) and 60% milliQ water as described in the prior study^12^ and administered by oral gavage, 25 mg/kg, once per day in a final volume of 100 μL. The control mice received the vehicle by oral gavage in the same dose and frequency. The mice were housed in modified barrier conditions, fed ad libitum (LabDiet 5053) in 12-hour light/ dark cycle at 21^0^C, 40-60% humidity. Both male and female mice were used for all the experiments described in this manuscript. All experiments were approved by MD Anderson Cancer Center IACUC.

### Cell culture

The KPC1 cells were cultured in 10% fetal bovine serum (FBS) RPMI (Corning, 10-040-CV) with 1% Penicillin-Streptomycin (PS) (Corning, 30-002-CI). The KPC2 and KPC3 cells were cultured in 10% FBS DMEM (Corning,10-017-CV) with 1%PS. All cells were tested and confirmed to be negative for mycoplasma using LookOut Mycoplasma PCR Detection Kit (Sigma-Aldrich, MP0035) prior to use in *in-vivo* experiments or analysis.

### Flowcytometry

PDAC tissues/ pancreas were minced with razor blades and digested in 1.5mg/mL of Type-1 Collagenase (Gibco, 17100017) in 10% Bovine serum albumin (Sigma-Aldrich, A9418) in 10mL of PBS at 37^0^C for 6-10 minutes in an incubator at 150rpm. The samples were subsequently washed in cRPMI (10% heat-inactivated FBS (HI-FBS)(Atlanta Biologicals, Atlanta, GA, USA), 1% PS, 1 mM sodium pyruvate (ThermoFischerScientific, 11360070), and 50 mM β-mercaptoethanol (LifeTechnologies, 31350010)) and centrifuged at 800g for 3 mins at room temperature 4^0^C. Cells were filtered using a 70μm strainer, washed and resuspended in FACS buffer (2% HI-FBS in PBS). Cells were stained in a final volume of 100μL with antibodies listed in **Table 1** diluted in 20% brilliant stain buffer (BD Bioscience,566349), Live/dead stain, 1:1000 (eBioscience, 65-0865-14), and anti-mouse CD16/32, 1:200 (TONBObiosciences, 40-0161) for 30 min on ice. Subsequently, cells were permeabilized (Foxp3/Transcription Factor Staining Buffer Set (eBioscience, 00-5523-00) as per manufacturer’s instructions and incubated with intra cellular antibodies for 30 mins following which the cells were fixed (BD Bioscience, 554655) and data were acquired using a Cytek Aurora and analyzed with FlowJo v10.7.1.

**Table 1:** Antibodies used for CD45^+^ immune cell analysis.

| **Antibodies/**<br>**Dyes** | **Fluorochrome** | **Catalog No.** |
| --- | --- | --- |
| GranzymeB | efluor 660 | Thermofisher 50-8898-82 |
| CD8 | BV650 | Biolegend 100742 |
| CD45 | Pacific Blue | Biolegend 103126 |
| Live dead<br>(FVD) | efluor 780 | eBiosciences 65-0865-14 |
| Ki67 | Alexa 488 | BD 558616 |
| CD4 | BUV395 | BD 565974 |
| FoxP3 | Percp Cy5.5 | eBiosciences 45-5773-80 |
| CD3 | Alexa 700 | eBiosciences 56-0032-82 |
| PD1 | BUV496 | BD 568597 |
| CD69 | PE | Invitrogen 12-0691-81 |
| TIM3/CD366 | PE/Dazzle (PE-TxR) | Biolegend 134013 |
| T-bet | PE-Cy7 | eBioscience 25-5825-80 |
| CD19 | Percp Cy5.5 | BD 551001 |

### Single cell RNA/ ATAC sequencing

PDAC tissues/ pancreas were minced and digested in 1.5mg/mL of Type-1 Collagenase in 10% Bovine serum albumin in 10mL at 37^0^C for 6-10 minutes in an incubator at 150 rpm. The samples were subsequently washed in cRPMI and stained with Live/dead (Fixable-viability dye (FVD)) 1:1000 and/or CD45 (1:100), CD16/32 (1:200) for 20 minutes and sorted for Live cells or CD45^+^ cells and submitted to the ATGC core for for single cell processing. Chromium Next GEM Single Cell 3 Reagent Kit v3.1 was used for scRNA seq. and Chromium Next GEM Single Cell Multiome ATAC + Gene Expression kit was used for scATAC seq. downstream processing as per the manufacturer’s protocol.

#### scRNA-seq Data Processing and Analysis

Raw FASTQ files were processed with Cell Ranger version 7.0.1 to generate count matrices. For aligning reads from various mouse models, we used the mm10 reference genome provided on the 10X Genomics website. Downstream analysis was performed in R (version 4.1.3) using the Seurat package (version 4.4.0) (Butler et al. 2018). We applied several Seurat functions to refine and interpret the data. First, to minimize the influence of noise and eliminate low-quality cells, we filtered out cells expressing an unusually low or high number of genes. Specifically, we determined filtering thresholds using R’s quantile function, retaining cells with gene counts between the 2.5th and 97.5th percentiles and discarding those outside this range. Additionally, cells exhibiting more than 5% mitochondrial counts were excluded from further analysis. Normalization and variance stabilization were achieved with the “Sctransform” function. We then reduced dimensionality via principal component analysis (PCA). The ‘FindNeighbors’ function identified nearest neighbors within the PCA space, and the optimal number of principal components was selected by examining outputs from both ‘JackStrawPlot’ and ‘ElbowPlot’. Cells were subsequently clustered using the ‘FindClusters’ function with an appropriate resolution setting, and the ‘RunUMAP’ function was employed to generate UMAP visualizations of the clusters. Finally, the ‘DoHeatmap’ function showcased the top 10 genes for each meta-cluster, while the ‘VinPlot’ function illustrated the distribution of gene expression probabilities across clusters. Gene imputation was carried out using the MAGIC (Markov Affinity-based Graph Imputation of Cells) algorithm^103^.

#### scATAC-seq Data Processing and Analysis

For scATAC-seq, raw data were initially processed with Cell Ranger ATAC version 2.1.0, which mapped reads to the mm10 genome and generated fragment files. These fragment files were then analyzed in R using the ArchR package (version 1.0.2). Quality control steps included removing cells with fewer than 4,000 unique fragments or a TSS enrichment score below three, and doublets were filtered out using default parameters. Cells that met these quality criteria were clustered using ArchR’s integration of the Seurat method at a resolution of 0.5. Dimensionality reduction and visualization were performed using UMAP, configured with 30 neighbors. Peak calling was conducted on aggregated coverage profiles for each identified cell population using MACS2 with its default settings. Additional analyses, such as motif enrichment and footprinting, were executed using the corresponding functions within ArchR under recommended settings.

### Immunostaining

Formalin-fixed paraffin-embedded (FFPE) tissues were sectioned (5 µm thickness), melted at 65°C for 1h and deparaffinized. Antigen retrieval was performed in Tris-EDTA buffer (pH 8.6) at 95°C for a total of 45 mins (20 mins at 95°C) using a steamer and cooled in an ice bath for 30 mins. Subsequently, using a Pap pen, a hydrophobic chamber was created surrounding the tissue, followed by washing in PBST (0.1% Tween 20) for 3 mins. The sections were then treated with 3% H_2_O_2_ for approximately 10-15 mins, followed by blocking with 1.5% bovine serum albumin in PBST for 1 hour at room temperature. Primary antibodies for phospho-p44/42 MAPK (Erk1/2) (Thr202/Tyr204) (Cell Signaling Technology, #4376S, 1:100), or CK19 (EP1580Y) (Abcam, #52625, 1:400) were applied to the sections and incubated for 3h at RT. Sections were then incubated with rabbit-on-rodent horseradish peroxidase polymer (BioCare Medical) for 15 minutes, followed by DAB (Life Technologies) incubation for 5–10 mins or until positive expression was seen. The sections were then counterstained with hematoxylin, dehydrated, and finally cover slipped. Representative histological images were assessed and collected using a Leica DM1000 LED microscope equipped with a DFC295 camera (Leica) and LAS version 4.4 software (Leica), with quantification analysis performed using ImageJ.

For CD8, CD19 and CD31, co-expression analysis, a Tyramide signal amplification (TSA) staining was conducted. 5 µm thickness sections were processed as described above until the blocking step. After blocking for 1h at RT using blocking buffer, the sections were further incubated with CD8 (D8A8Y) (Cell Signaling Technology, #85336S, 1:200) primary antibody for 3h at RT. The slides were then washed and further incubated with rabbit-on-rodent HRP polymer (BioCare Medical) for 15 mins, followed by a 30-min incubation with Opal 690 (Akoya Biosciences, 1:100) followed by a second antigen retrieval step. Next, for CD19 staining, sections incubated in blocking buffer, followed by incubation in CD19(D4V4B) (Cell Signaling Technology, #90176S, 1:100) for 3h at RT. The sections were then incubated in rabbit on rodent HRP polymer for 15 mins, followed by 30 min incubation in Opal 520 (Akoya Biosciences, 1:100). Next, following antigen retrieval, the sections were incubated in CD31 primary antibody (D8V9E) (Cell Signaling Technology, #77699S), subsequent incubation in rabbit-on-rodent HRP polymer and Opal 570 (Akoya Biosciences, 1:100) followed by a final antigen retrieval step and cover slipping with Fluoroshield with DAPI (Sigma Aldrich, F6057). Similarly, a TSA for CD21 and CD21 expression was performed (Reagent details in Table 2). All TSA images were observed and collected using a Zeiss LSM800 confocal microscope and quantified using Image J.

**Table 2:**
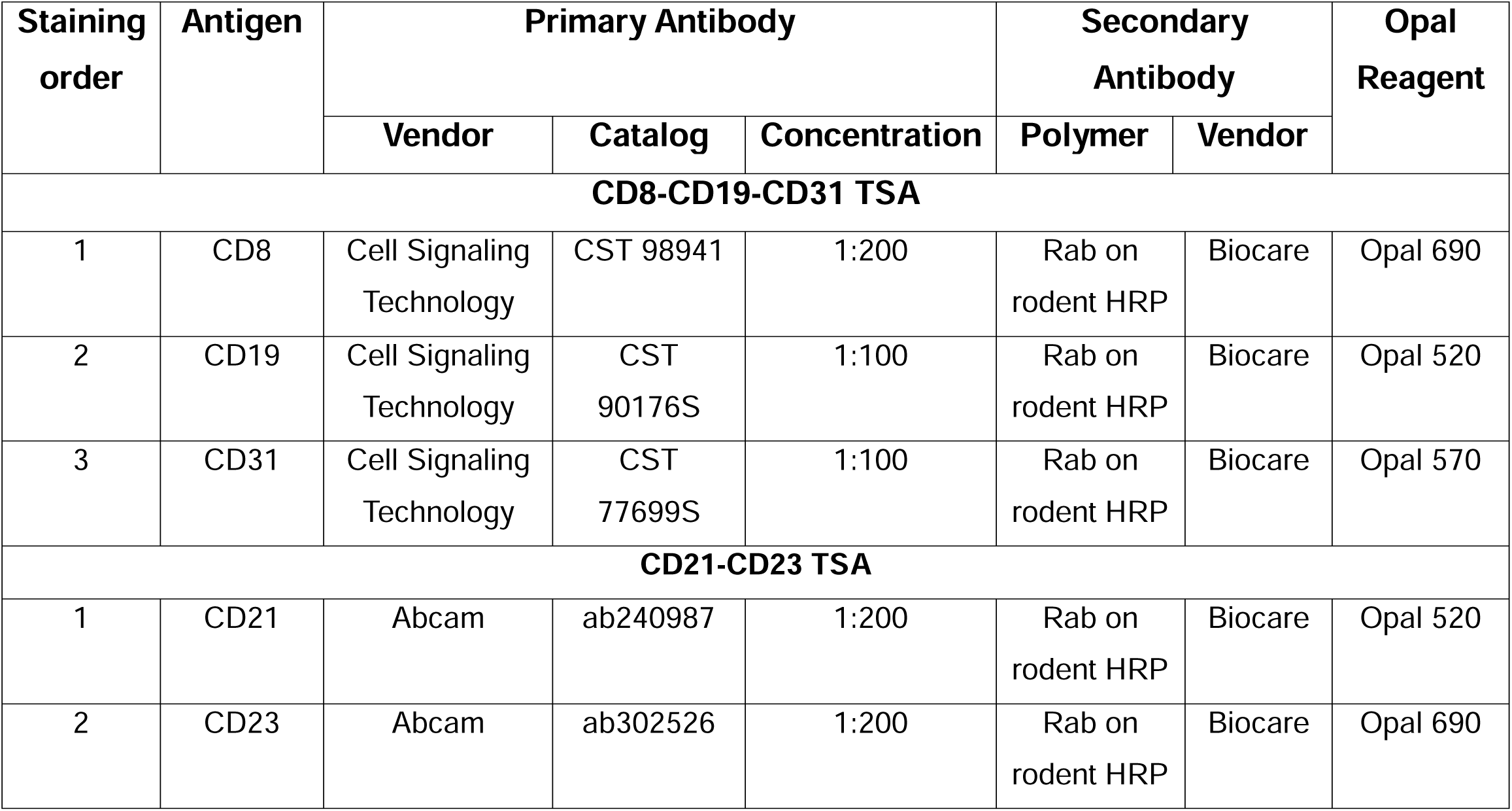
Antibodies and reagents used for TSA staining.

### Statistical analysis

Figures in the manuscript were generated using Graphpad prism 9 and R-studio. Normality of distribution of datasets was assessed using Shapiro-Wilk test. For comparisons involving two groups, unpaired-T test (datasets with normal distribution) or Mann-Whitney test (datasets with non-normal distribution) was used. For comparisons of multiple groups, one-way ANOVA (datasets with normal distribution) or Kruskal-Wallis test (datasets with non-normal distribution) was used. Log-rank test was used for survival analysis and two-way ANOVA was used for histological comparisons. *P* values throughout the manuscript represent \**P* < 0.05, \*\**P*< 0.01, \*\*\**P* < 0.001, \*\*\*\**P* < 0.0001; *ns*, not significant.

## Acknowledgements

We thank Navid Sobhani for assistance with single cell RNA Seq. of the KPC1 tumors and Haoqiang Ying for kindly providing the HY19636 PKC cells. We thank Barbara Moreno-Diaz and Austin Dellafosse for assistance with tissue processing and sectioning respectively. We thank Michelle Kirtley, Sujuan Yang, Patience Kelly and Kathleen McAndrews for help with mouse colony. We thank David Pollock from the Advanced Technology Genomics core (ATGC), MD Anderson Cancer Center for assistance with single cell sequencing experiments, Karen Ramirez and Veena Papanna from the Advanced Cytometry and Sorting Facility (ACSF), MD Anderson Cancer Center for cell sorting.

## Funding

Research in the Kalluri lab is funded by Break Through Cancer and the Lustgarten Foundation for Pancreatic Cancer Research. K.K.M. is supported by The Ergon Foundation Post-Doctoral Fellowship. The Kalluri laboratory PDAC work is supported by NCI P01 CA117969. A.M. is supported by the Sheikh Khalifa bin Zayed Foundation. A.P. is supported by POLISH-U.S. FULBRIGHT COMMISSION, Junior Research Award 2023-24, Grant Authorization Form No. PL/2023/19/JR. The ATGC core is supported by CA0126672 and NIH 1S10OD024977-01 award.

## Conflict of interest

A.M. is a consultant for Tezcat Biosciences and is named on a patent that has been licensed to Thrive Earlier Detection (an Exact Sciences company).

## Author contributions

K.K.M. and R.K. conceived, designed the project and wrote the manuscript. K.K.M., A.M., D.S.H., S.S., H.S., B.L., T.P.H. reviewed and edited the manuscript. K.K.M., A.S.M, A.B., S.V.K., A.M.S., S.J.M., K.A., H.S., performed experiments. T.P.H. generated and provided small molecule inhibitors. K.K.M, B.L., and A.K.P. analyzed the data. K.K.M., A.S.M. performed and supervised *in-vivo* studies. K.K.M., A.S.M., A.B., S.J.M., A.M.S. assisted with tissue processing and staining. K.K.M., K.A., and S.V.K. performed single cell sequencing experiments. K.K.M., B.L. and A.K.P. generated figures and analyzed single cell sequencing data.

---

**Supplementary Figure 1:**
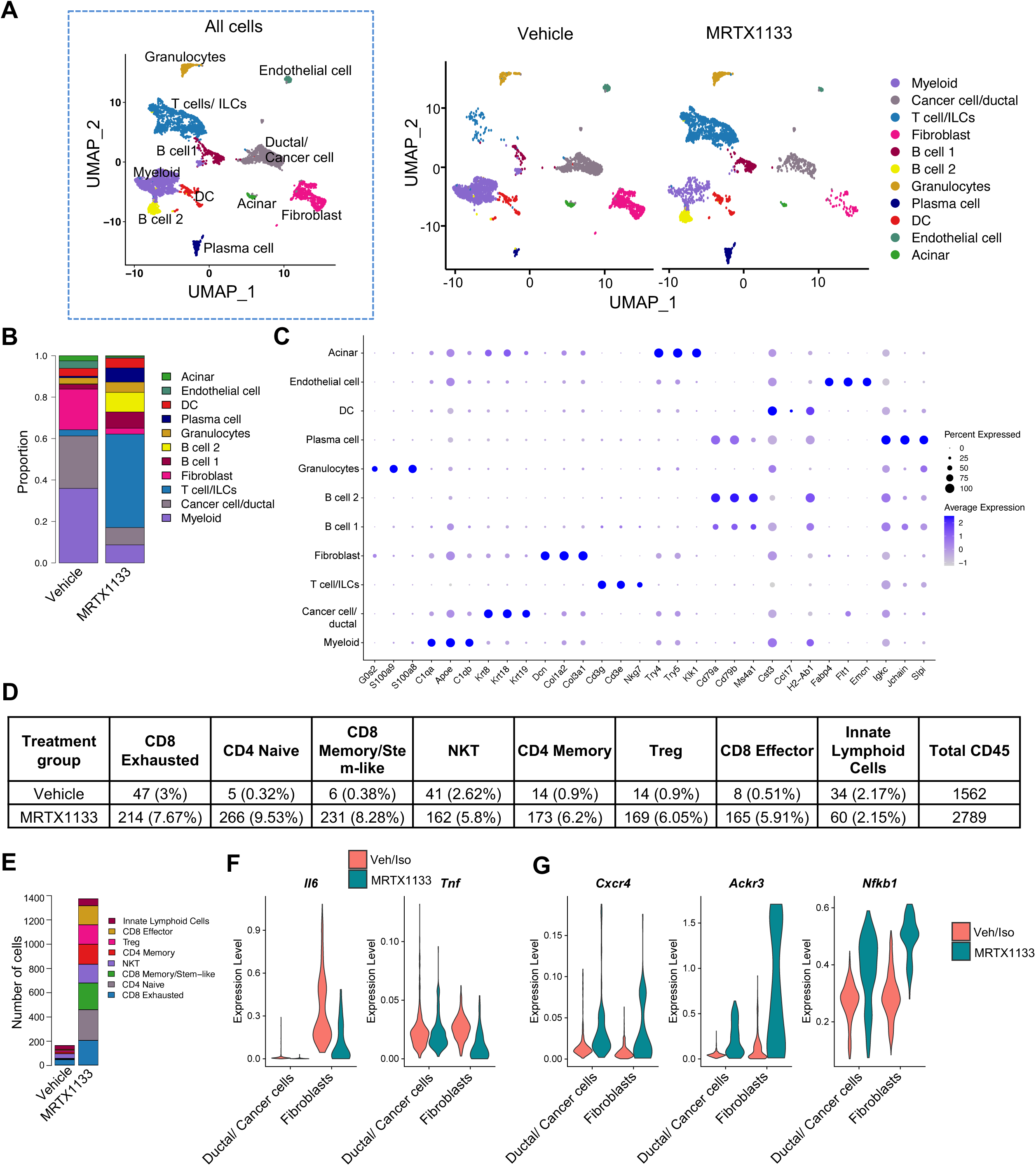
scRNA seq. analysis of KPC1 PDAC treated with MRTX1133. Related to Fig. 1. **(A)** UMAP of all cell types combined (left) and UMAP of all cells in Vehicle and MRTX1133 treated (right) KPC1 PDAC determined by scRNA seq. analysis (n=3 tumors/ group). **(B)** Relative abundance of all cell types in KPC1 PDAC determined by scRNA seq. **(C)** Dot plots of genes used to define distinct cell types by scRNA seq. **(D-E)** Number of T cells as table **(D)**, and bar graph **(E)** in KPC1 PDAC determined by scRNA seq. from Fig. 1E. **(F-G)** Violin plots for *Il6*, *Tnf* **(F)** and *Cxcr4, Ackr3*, and *Nfkb1* **(G)** in ductal/ cancer cells and fibroblasts of KPC1 mice determined by scRNA seq.. Data are presented as mean in **B, E** and as violin plots in **F** and **G**. Note: UMAP from Fig. 1B is generated from the CD45^+^ cells expressing *Ptprc* in S1A.

**Supplementary Figure 2:**
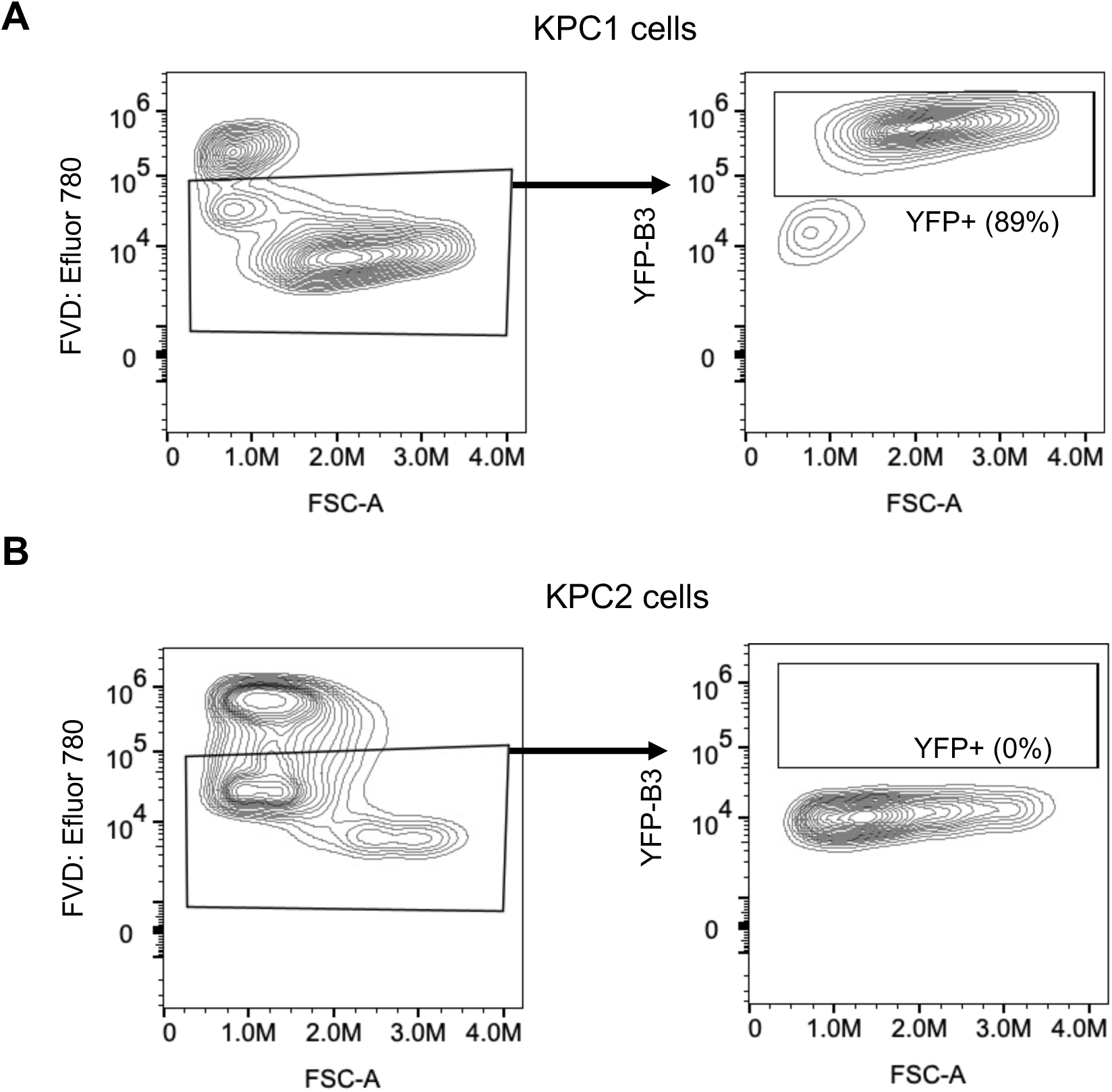
YFP expression in KPC1 and KPC2 cell lines. Related to Fig. 2. **(A-B)** Flow cytometry analysis with gating strategy for endogenous YFP expression in KPC1 **(A)**, and KPC2 **(B)** cells as a percentage of live cells.

**Supplementary Figure 3:**
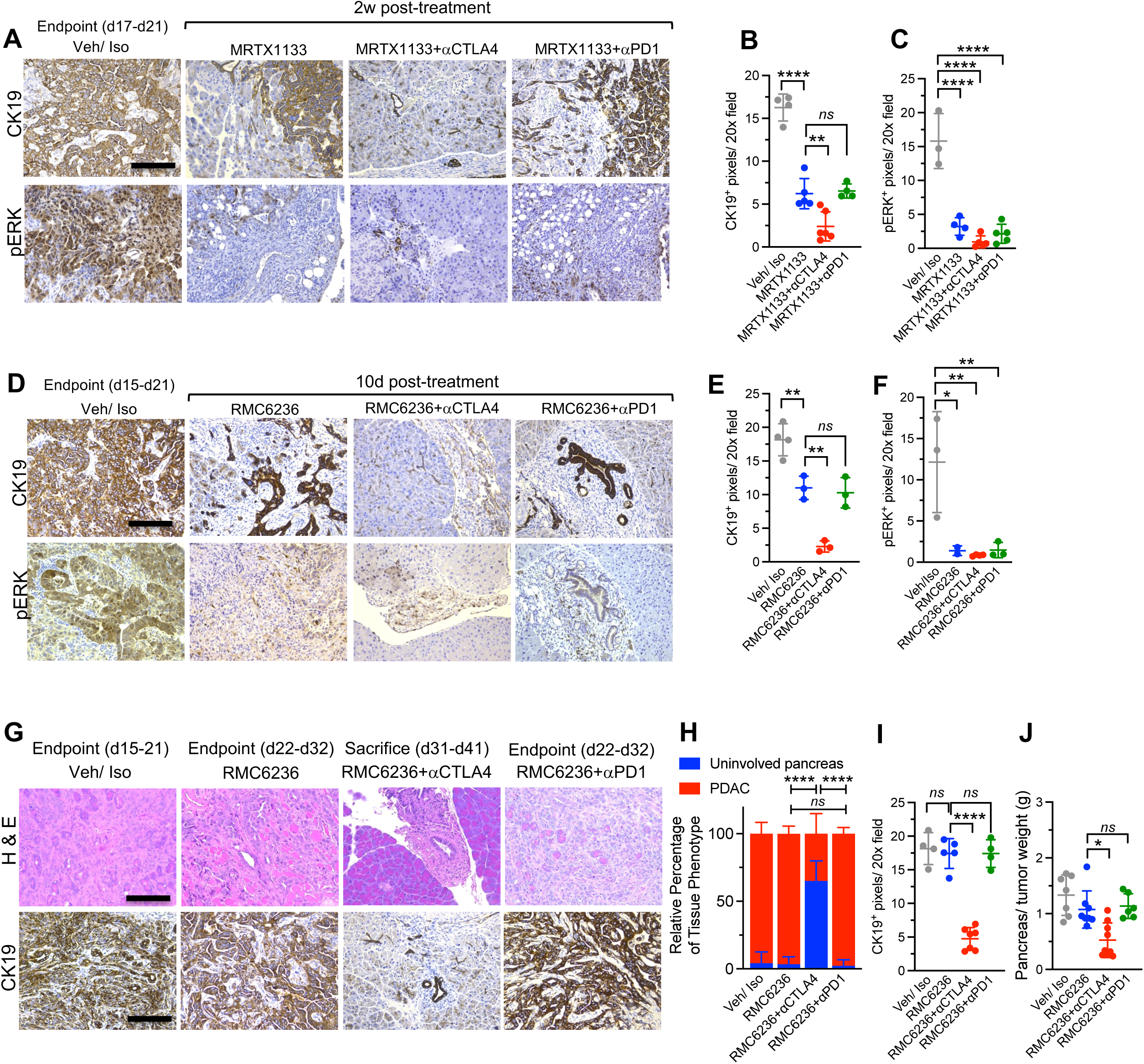
Additional histological and immunostaining analysis of KPC2 PDAC treated with MRTX1133, RMC6236 in combination with checkpoint immunotherapy. Related to Fig. 2. **(A-C)** Representative CK19 IHC (top) and pERK IHC (bottom) in age-matched KPC2 mice treated with Veh/Iso, MRTX1133, MRTX1133+αCTLA4, MRTX1133+αPD1 **(A)**, with quantification of CK19^+^ **(B)** and pERK^+^ **(C)** pixels per 20x field (n=3-5 tumors/ group). **(D-F)** Representative CK19 IHC (top) and pERK IHC (bottom) in age-matched KPC2 mice treated with Veh/Iso, RMC6236, RMC6236+αCTLA4, RMC6236+αPD1 **(D)**, with quantification of CK19^+^ **(E)** and pERK^+^ **(F)** pixels per 20x field (n=3-5 tumors/ group). **(G-I)** Representative H&E (top) and CK19 IHC (bottom) in endpoint KPC2 mice treated with Veh/Iso, RMC6236, RMC6236+αCTLA4, RMC6236+αPD1 **(G)**, with quantification of histology **(H)** and CK19^+^ pixels **(I)** pixels per 20x field (n=4-6 tumors/ group). Note: Same Iso/Veh groups were used in D, G. Data are presented as mean ± SD in **B, C, E, F, H, I** and **J**. Significance was determined by one-way ANOVA with Sidak’s multiple comparisons test in **B, C, E, F** and **I**, Kruskal-Wallis with Dunn’s multiple comparisons test in **J**, and two-way ANOVA in **H.** Scale bars-100mm. \**P* < 0.05, \*\**P*< 0.01, \*\*\**P* < 0.001, \*\*\*\**P* < 0.0001; *ns*, not significant.

**Supplementary Figure 4:**
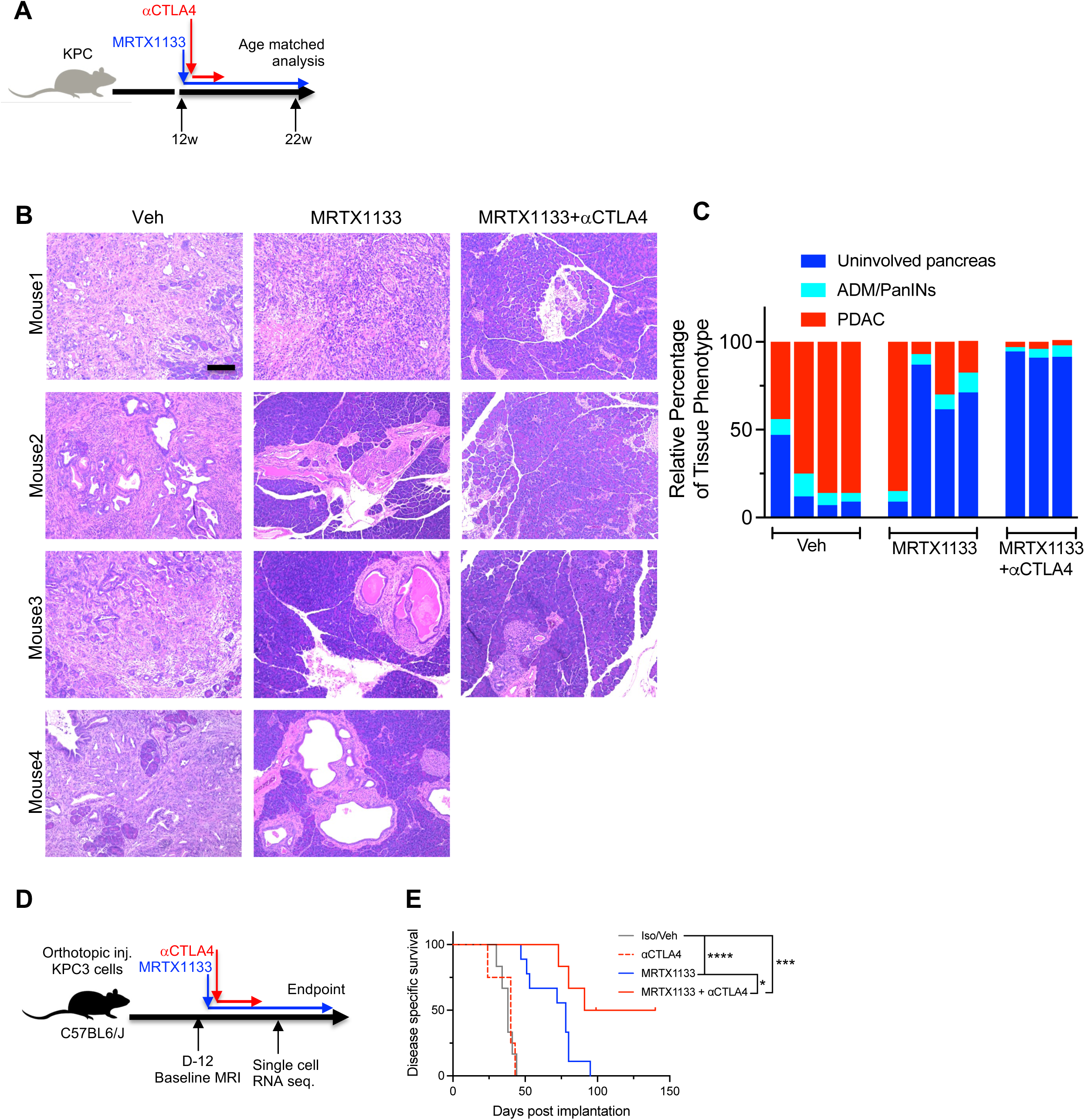
Additional analysis of autochthonous KPC tumors and KPC3 tumors. Related to Fig. 2. **(A)** Schematic representation of treatment of KPC GEMs with MRTX1133 and αCTLA4. **(B-C)** Representative H&E images (B) with quantification of relative percentage of histological phenotype **(C)** of individual KPC mice (age matched) treated with Veh/Iso, MRTX1133, and MRTX1133+αCTLA4 (n=3-4 mice/ group). **(D)** Schematic representation of orthotopic injection and MRTX1133 and αCTLA4 treatments of KPC3 mice. **(E)** Kaplan-Meier survival curves of Veh/Iso (n=6), αCTLA4 (n=4), MRTX1133 (n=9), MRTX1133+αCTLA4 (n=6) treated KPC3 mice. <u>Note:</u> Some of the mice in the KPC3 survival curve has been used earlier^24^. Data are presented as mean in **C**. Significance was determined by log-rank test in **E.** Scale bars-100mm. \**P* < 0.05, \*\*\**P* < 0.001, \*\*\*\**P* < 0.0001.

**Supplementary Figure 5:**
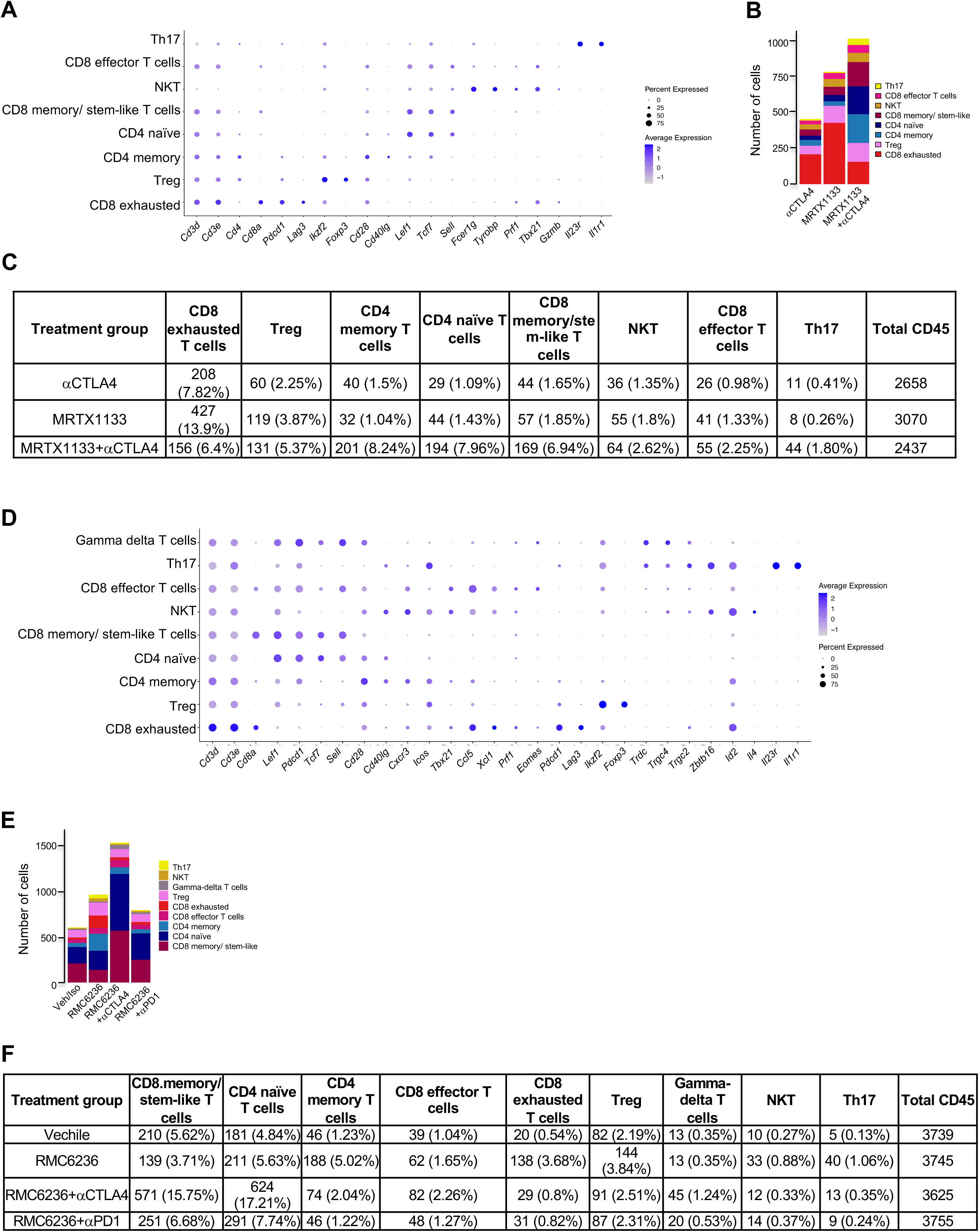
Additional scRNA seq. analysis of MRTX1133, RMC6236 treated PDAC. Related to Fig. 3. **(A)** Dot plots of genes used to define T cell subtypes in MRTX1133 treated KPC3 determined by scRNA seq. analysis. **(B-C)** Number of T cells as bar-graph **(B)**, and table **(C)** in KPC3 PDAC determined by scRNA seq. from Fig. 3A-B. **(D)** Dot plots of genes used to define T cell subtypes in RMC6236 treated KPC2 determined by scRNA seq. analysis. **(E-F)** Number of T cells as bar-graph **(E)**, and table **(F)** in KPC2 PDAC determined by scRNA seq. from Fig. 3C-D.

**Supplementary Figure 6:**
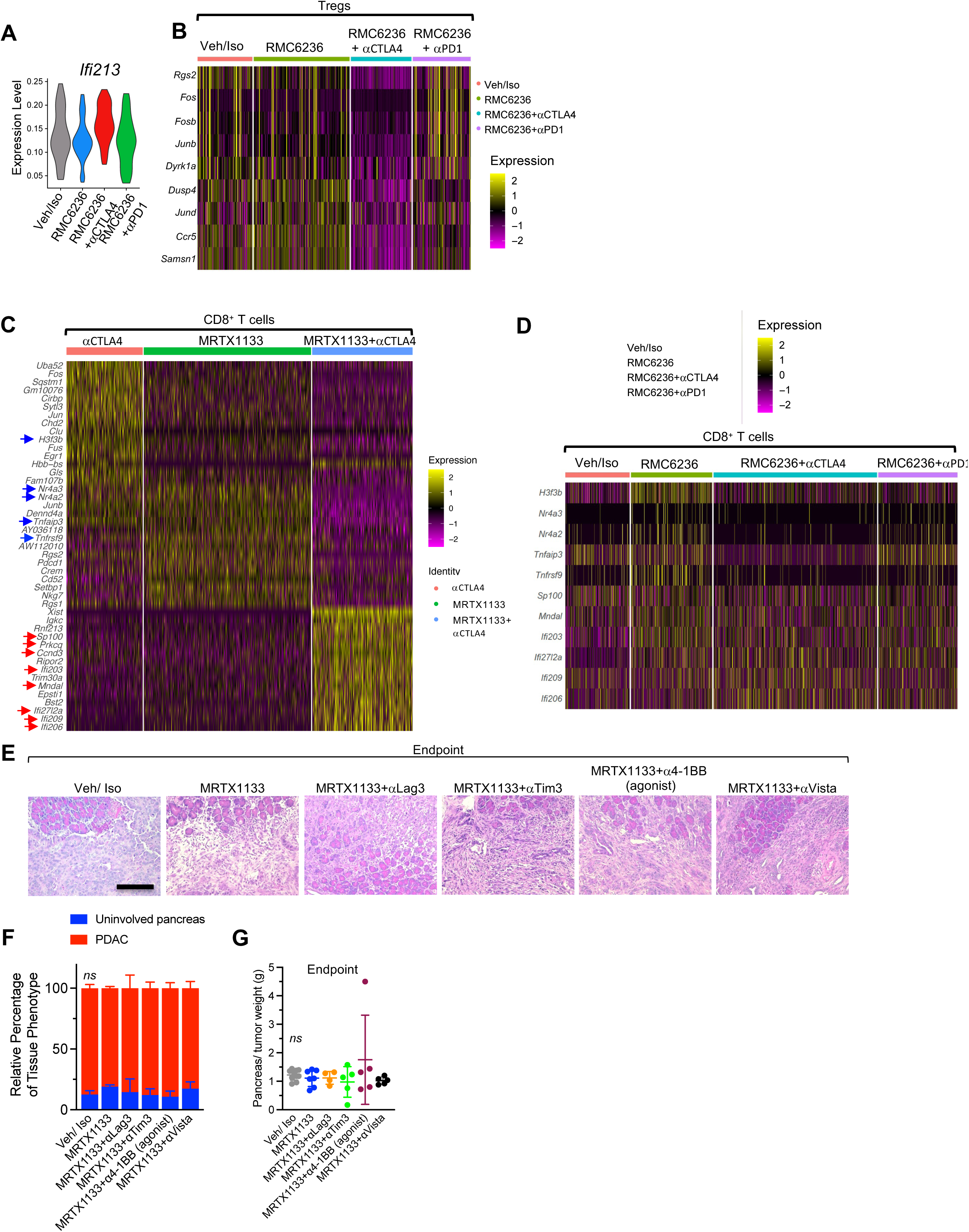
Additional scRNA seq. and histological analysis of Kras* targeting in combination with checkpoint immunotherapy in PDAC. Related to Fig. 3. **(A)** Violin plots for *Ifi213* in tumor infiltrating Tregs of RMC6236 treated KPC2 mice with aCTLA4/aPD1. **(B)** Heatmap of differentially regulated genes in the tumor infiltrating Tregs of Iso/Veh, RMC6236, MRTX1133+αCTLA4, RMC6236+ αPD1 treated KPC2 PDAC. **(C)** Heatmap of differentially regulated genes in the tumor infiltrating CD8^+^ T cells of αCTLA4, MRTX1133, MRTX1133+αCTLA4 treated KPC3 PDAC. (Red arrow indicates upregulated genes and blue arrow indicates downregulated genes in the MRTX1133+αCTLA4 group compared to the other groups). **(D)** Heatmap of differentially regulated genes in the tumor infiltrating CD8^+^ T cells of Iso/Veh, RMC6236, MRTX1133+αCTLA4, RMC6236+ αPD1 treated KPC2 PDAC. **(E-F)** Representative H&E in endpoint KPC2 mice treated with Veh/Iso, MRTX1133, MRTX1133+αLag3, MRTX1133+αTim3, MRTX1133+αVista, and MRTX1133+α4-1BB agonist antibody **(E)**, with quantification of histology **(F)** (n=2-5 tumors/ group). **(G)** Pancreas/ tumor weights of age-matched KPC2 mice with indicated treatment groups (n=4-10/ group). <u>Note:</u> Veh/ Iso group used in E-G here is repeated from Fig. 2F. Data are represented as violin plots in **A** and mean ± SD in **F** and **G**. Significance was determined by two-way ANOVA in **F** and one-way ANOVA with Dunnett’s multiple comparisons test in **G**. Scale bars-100mm. *ns*, not significant.

**Supplementary Figure 7:**
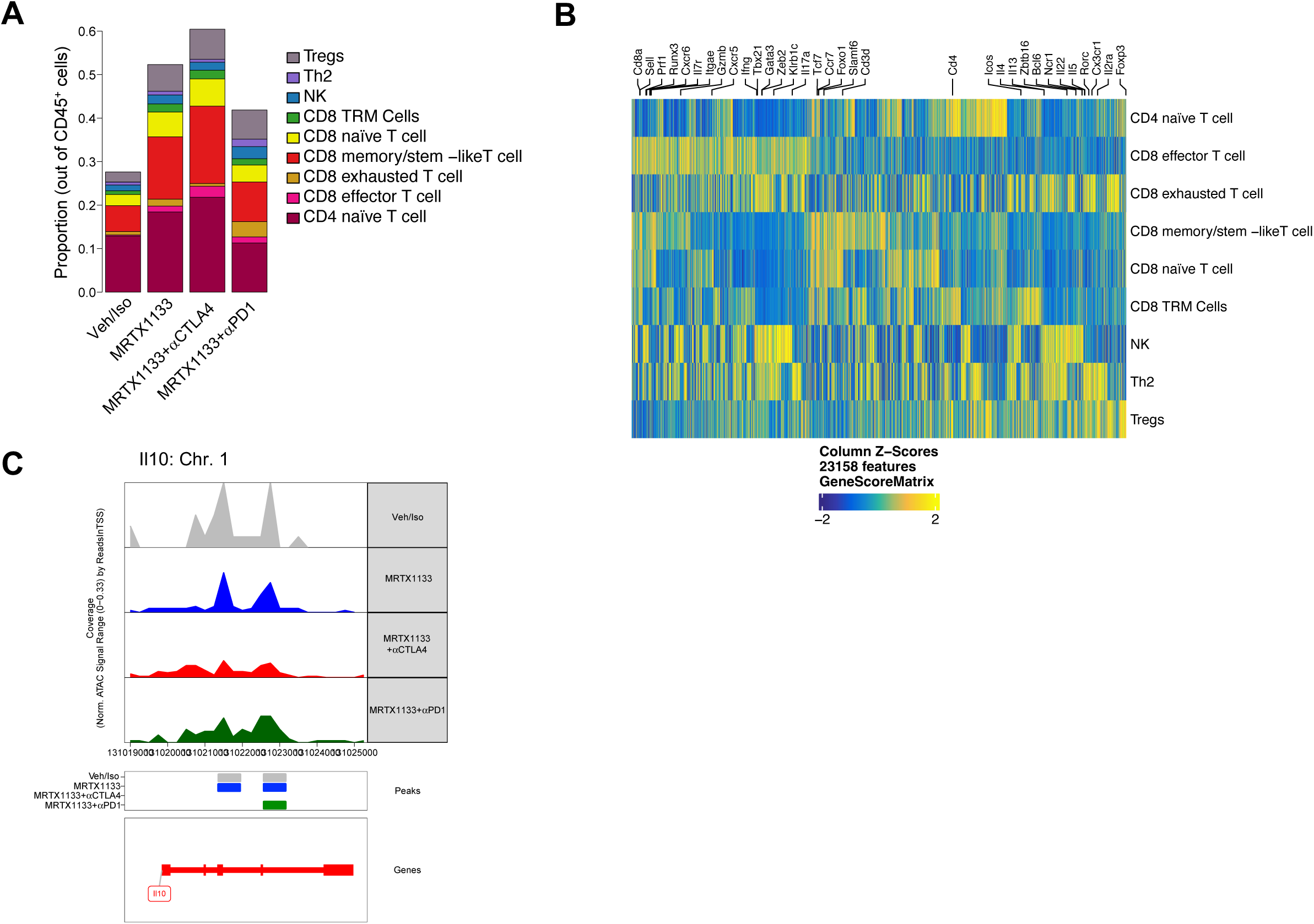
Additional scATAC seq. of tumor infiltrating Tregs in MRTX1133 and checkpoint immunotherapy treated PDAC. Related to Fig. 4. **(A)** Relative frequencies of T cell subsets (as % of CD45^+^ cells) in Veh/Iso, MRTX1133, MRTX1133+αCTLA4 and MRTX1133+αPD1 treated mice. n=3 tumors/ group. **(B)** Marker genes calculated based on gene scores to define T cell clusters. **(C)** Genome browser tracks of Tregs around the Il10 gene locus in Veh/Iso, MRTX1133, MRTX1133+αCTLA4, MRTX1133+αPD1 treated KPC2 PDAC. Data are presented as mean in **(A)**.

**Supplementary Figure 8:**
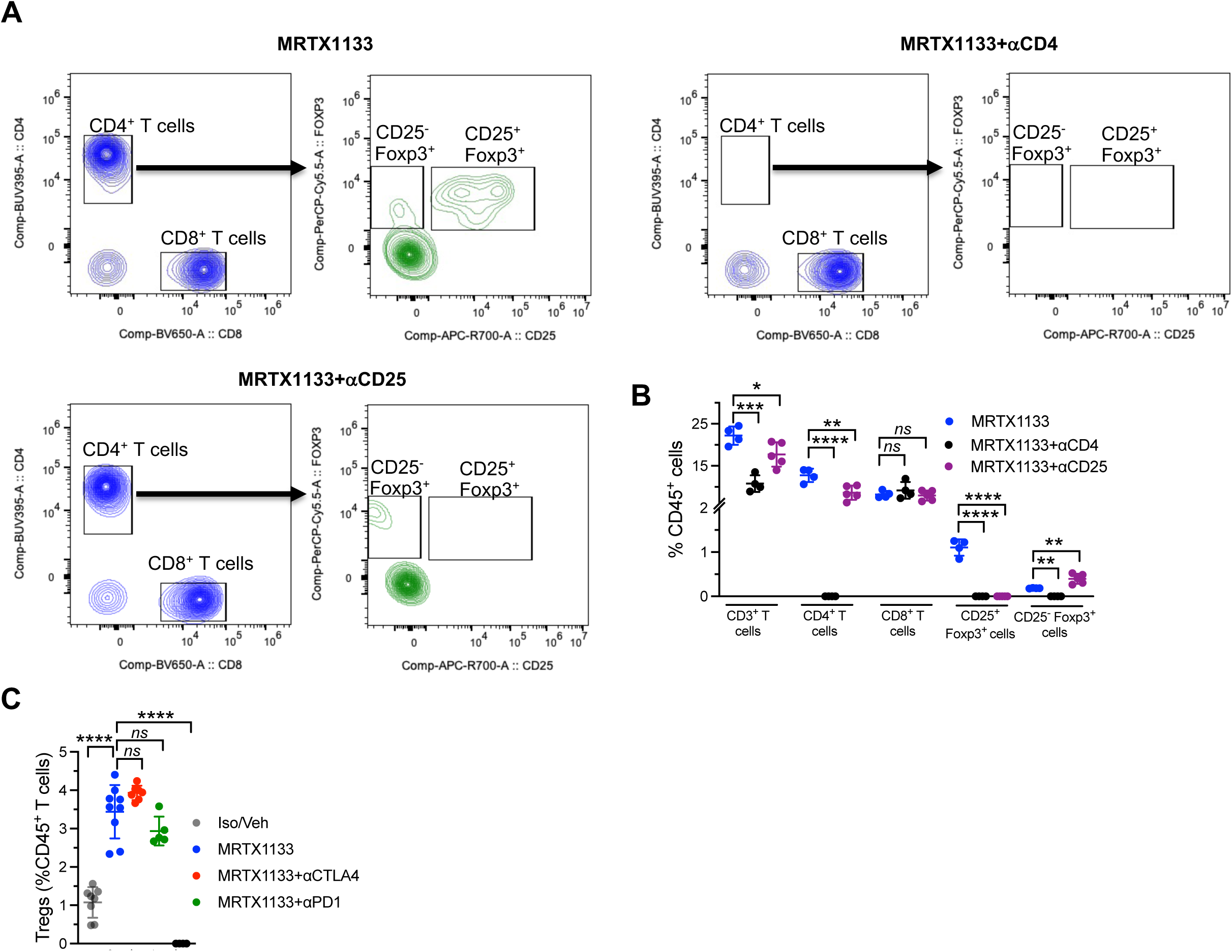
Additional flowcytometry analysis of T cell populations from Fig. 5. **(A-B)** Representative images of gating strategy in the peripheral blood of mice treated with MRTX1133, aCD4 and aCD25 depletion antibodies **(A)**, with quantification of frequencies of indicated cell populations **(B)** (n=4-5 mice/ group). **(C)** Immunophenotyping analysis of frequencies of CD4^+^Foxp3^+^ measured as a percentage of CD45^+^ cells in indicated treatment groups (n=4-9 tumors/ group). Data are presented as mean ± SD in B and C. Significance was determined by one-way ANOVA with Dunnett’s multiple comparisons test in B and C. \**P* < 0.05, \*\**P*< 0.01, \*\*\*\**P* < 0.0001; *ns*, not significant.

**Supplementary Figure 9:**
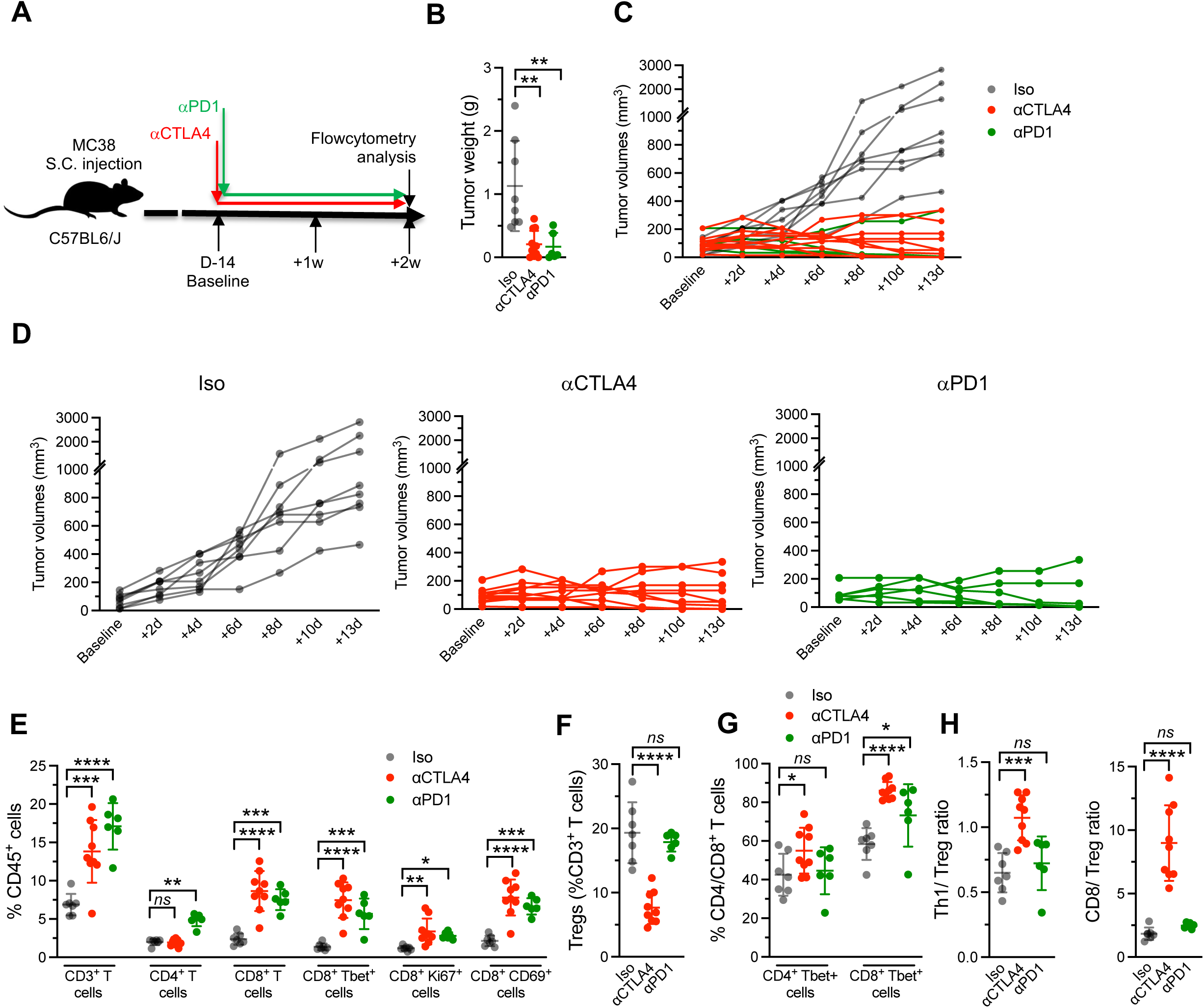
Tumor growth kinetics and flowcytometry analysis of MC38 tumors treated with checkpoint blockade immunotherapy. Related to Fig. 5. **(A)** Schematic representation of subcutaneous injection and aCTLA4 and aPD1 treatments of MC38 mice. **(B)** Tumor weights of age-matched MC38 mice treated with Iso (n=8), αCTLA4 (n=11), αPD1 (n=6). **(C)** Growth kinetics of Iso (n=8), αCTLA4 (n=11), αPD1 (n=6) treated MC38 tumors. **(D)** Growth kinetics of Iso (n=8), αCTLA4 (n=11), αPD1 (n=6) treated MC38 tumors from Supplementary Fig. S9C visualized as separate groups. **(E-H)** Immunophenotyping analysis of MC38 tumors of indicated groups. CD3^+^(T cells); CD3^+^CD4^+^ (CD4^+^ T cells); CD3^+^CD8^+^ (CD8^+^ T cells), CD8^+^ T-bet^+^ cells, CD8^+^ Ki67^+^ cells and CD8^+^ CD69^+^ cells were measured as a percentage of CD45^+^ cells **(E)**; CD4^+^Foxp3^+^ were measured as a percentage of CD3^+^ cells **(F)**; CD4^+^T-bet^+^ (Th1) and CD8^+^T-bet^+^ measured as a percentage of CD3^+^ cells **(G)**. CD8/Treg ratio and Th1/Treg ratio **(H)**. n=6-9 tumors/ group. Data are presented as mean ± SD in **B, C, E, F, G** and **H** and as individual tumor volume measurements in **C, D**. Significance was determined by Kruskal-Wallis with Dunn’s multiple comparisons test for CD4 comparisons in **E**, one-way ANOVA with Dunnett’s multiple comparisons test in **B, F, G** and **H** and other panels in **E**. \**P* < 0.05, \*\**P*< 0.01, \*\*\**P* < 0.001, \*\*\*\**P* < 0.0001; *ns*, not significant.

**Supplementary Figure 10:**
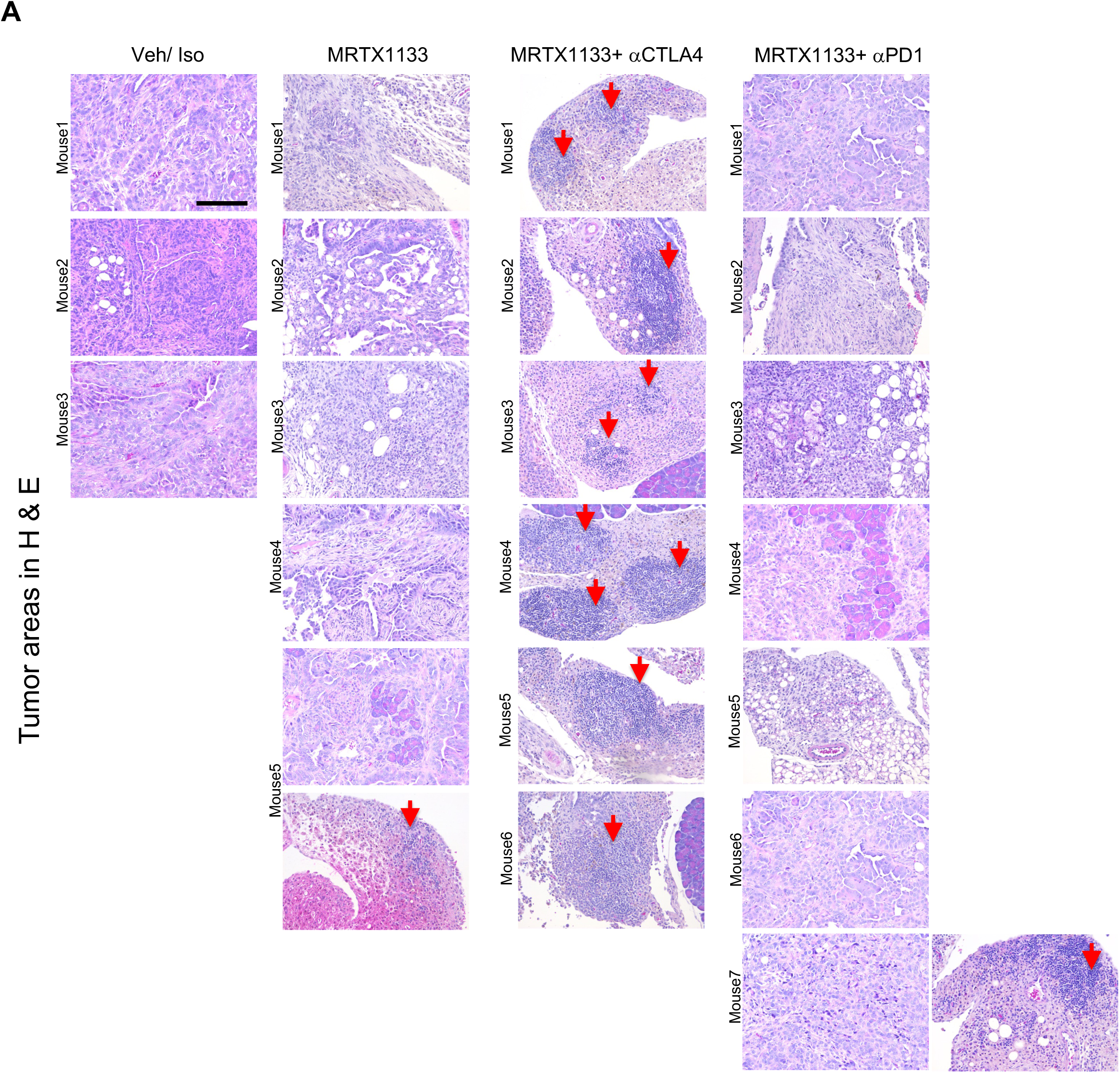
Additional histological analysis of KPC2 PDAC treated with MRTX1133 and checkpoint immunotherapy. Related to Fig. 6. **(A)** Representative individual H&E images of tumoral areas with TLS lesions in age-matched Veh/Iso, MRTX1133, MRTX1133+αCTLA4, MRTX1133+αPD1 treated KPC2 PDAC. Scale bars-100μm.

**Supplementary Figure 11:**
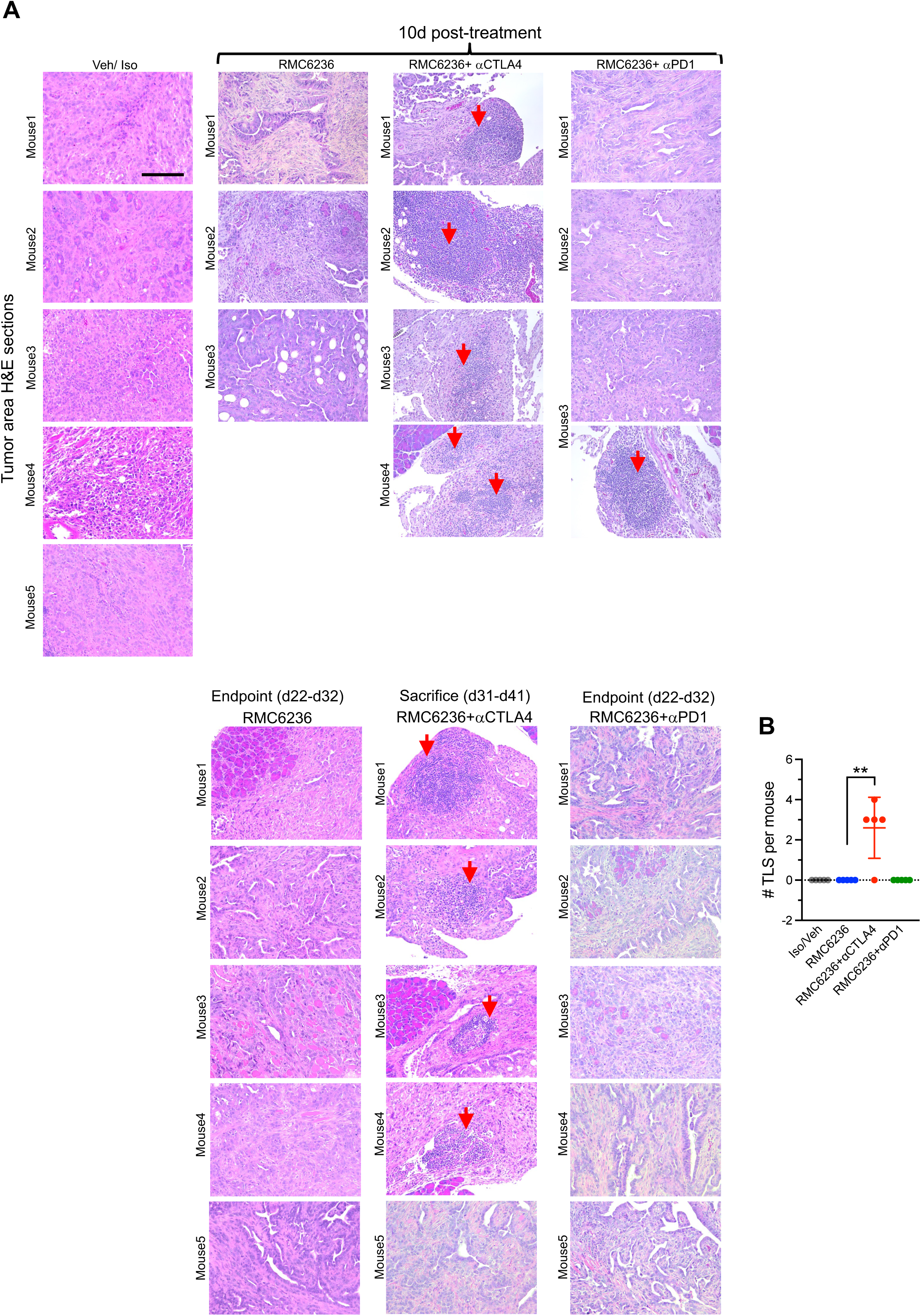
Additional histological analysis of KPC2 PDAC treated with RMC6236 and checkpoint immunotherapy. Related to Fig. 6. **(A-B)** Representative individual H&E images of tumoral areas with TLS lesions in age-matched (top panel) and endpoint (bottom panel) Veh/Iso, RMC6236, RMC6236+αCTLA4, RMC6236+αPD1 treated KPC2 PDAC **(A)**, with quantification of TLS in endpoint tumors **(B)**. Data are represented as mean ± SD. Significance was determined by unpaired T-test. Scale bars-100mm.\*\**P*< 0.01.

**Supplementary Figure 12:**
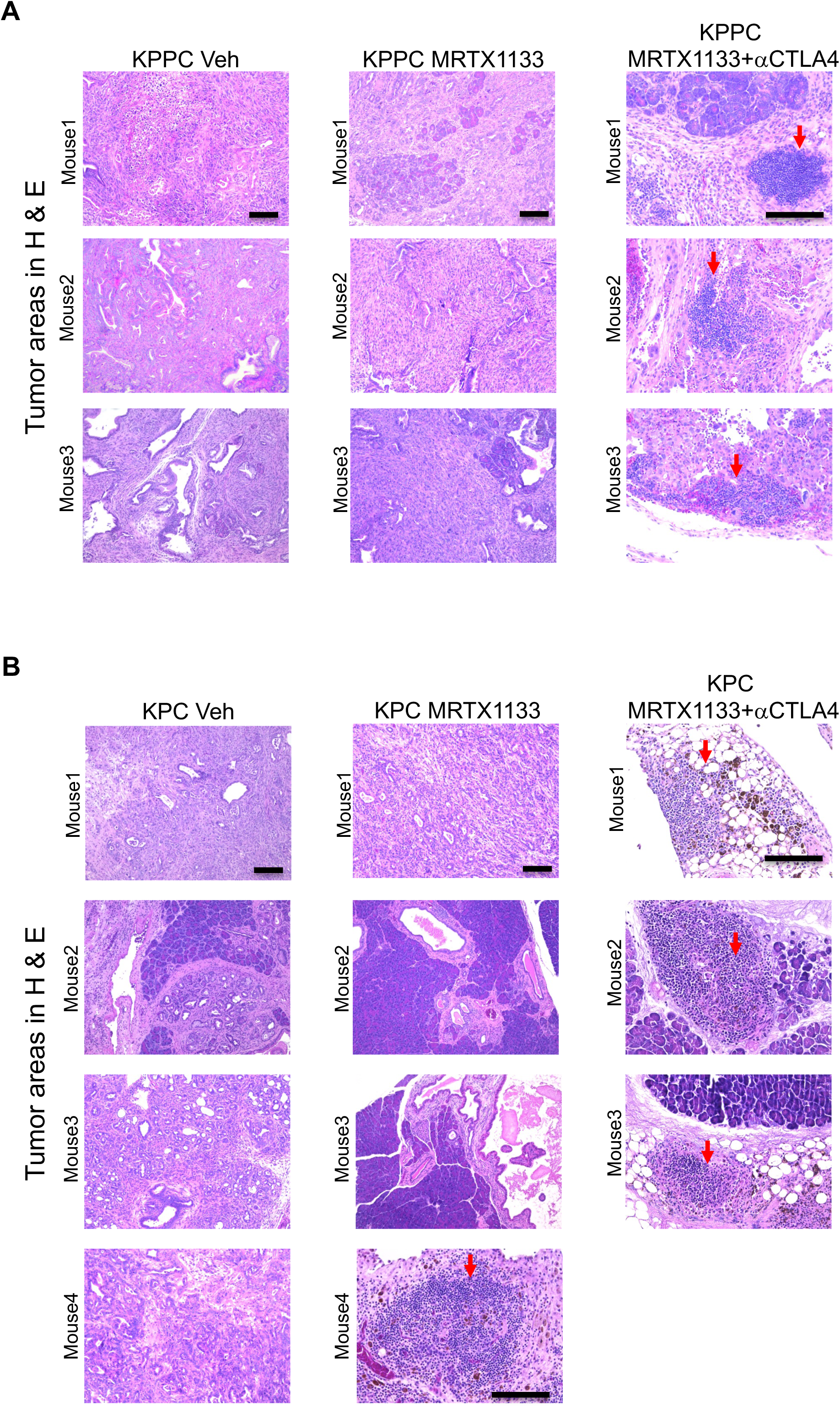
Additional histological analysis of KPC and KPPC GEMs treated with MRTX1133 and checkpoint immunotherapy. Related to Fig. 6. **(A)** Representative individual H&E images of tumoral areas with TLS lesions in age-matched Veh/Iso, MRTX1133, MRTX1133+αCTLA4 treated KPPC mice (n=3 mice/ group). **(B)** Representative individual H&E images of tumoral areas with TLS lesions in age-matched Veh/Iso, MRTX1133, MRTX1133+αCTLA4 treated KPC mice (n=3-4 mice/ group). Scale bars-100μm.

**Supplementary Figure 13:**
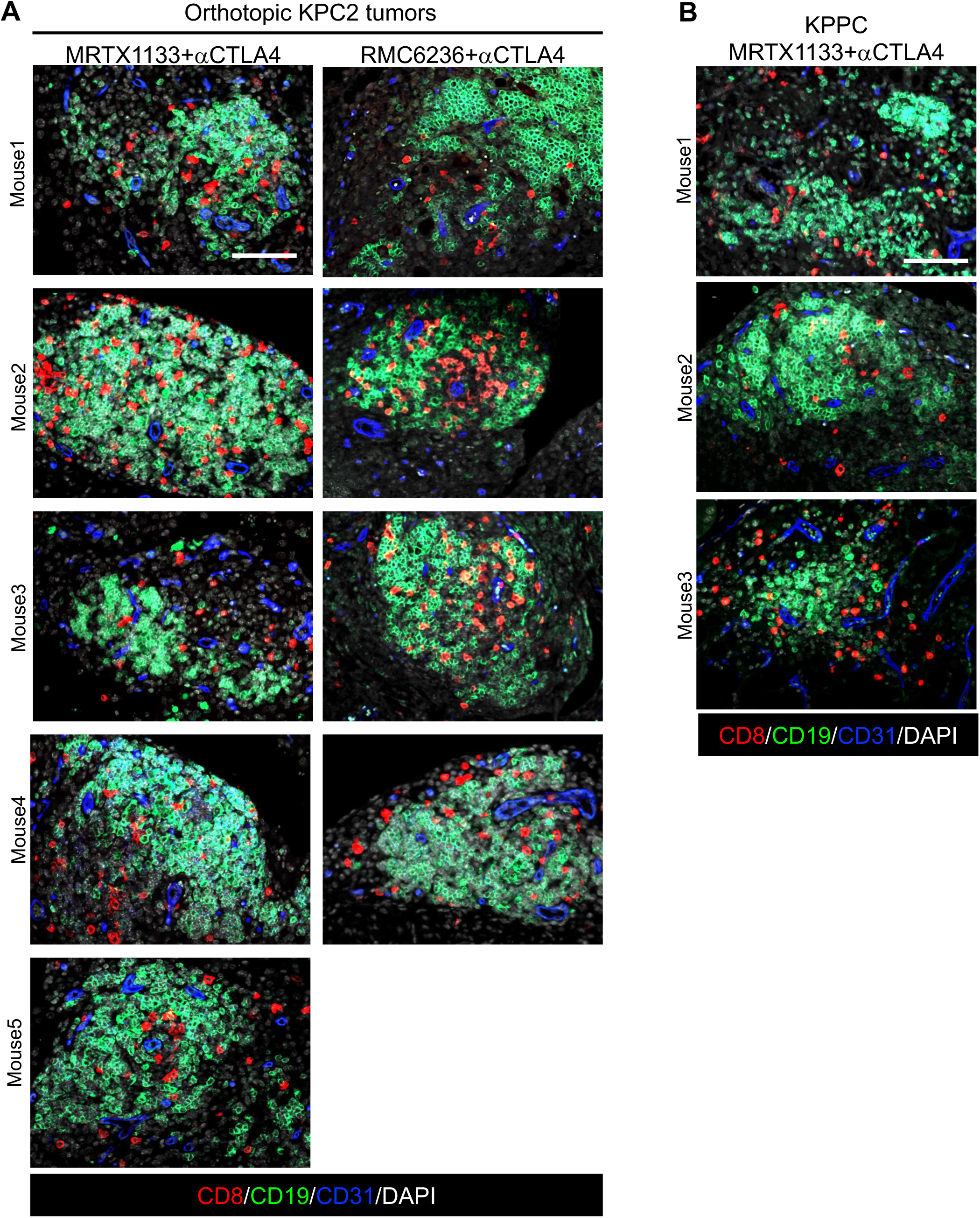
Additional immunostaining analysis of tertiary lymphoid structures from individual KPC2 and KPPC tumors. Related to Fig. 6. **(A)** Representative individual CD8-CD19-CD31 immunostaining images showing TLS in orthotopic KPC2 mice treated with MRTX1133/ RMC6236 and anti-CTLAA4. **(B)** Representative individual CD8-CD19-CD31 immunostaining images showing TLS in autochthonous KPPC mice treated with MRTX1133 and anti-CTLAA4. Scale bars-100mm.

**Supplementary Figure 14:**
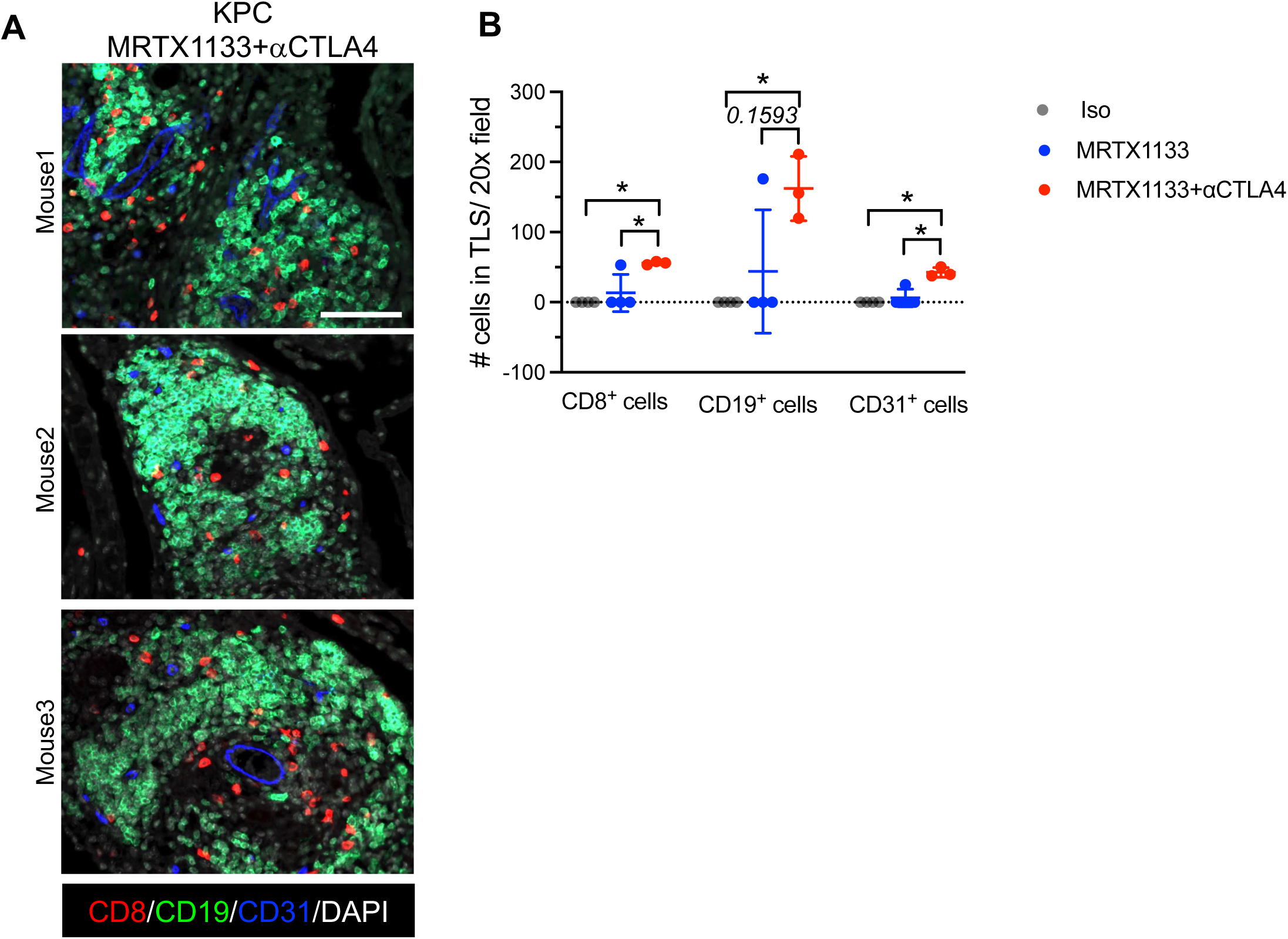
Additional immunostaining analysis of tertiary lymphoid structures from individual KPC tumors. Related to Fig. 6. **(A-B)** Representative individual CD8-CD19-CD31 immunostaining images **(A)**, with quantification of CD8, CD19 and CD31^+^ cells in TLS **(B)** of autochthonous KPC mice treated with MRTX1133 and anti-CTLAA4. Data are presented as mean ± SD in **B**. Significance was determined by Kruskal-Wallis with Dunn’s multiple comparisons test in **B.** \**P* < 0.05. Scale bars-100μm.

**Supplementary Figure 15:**
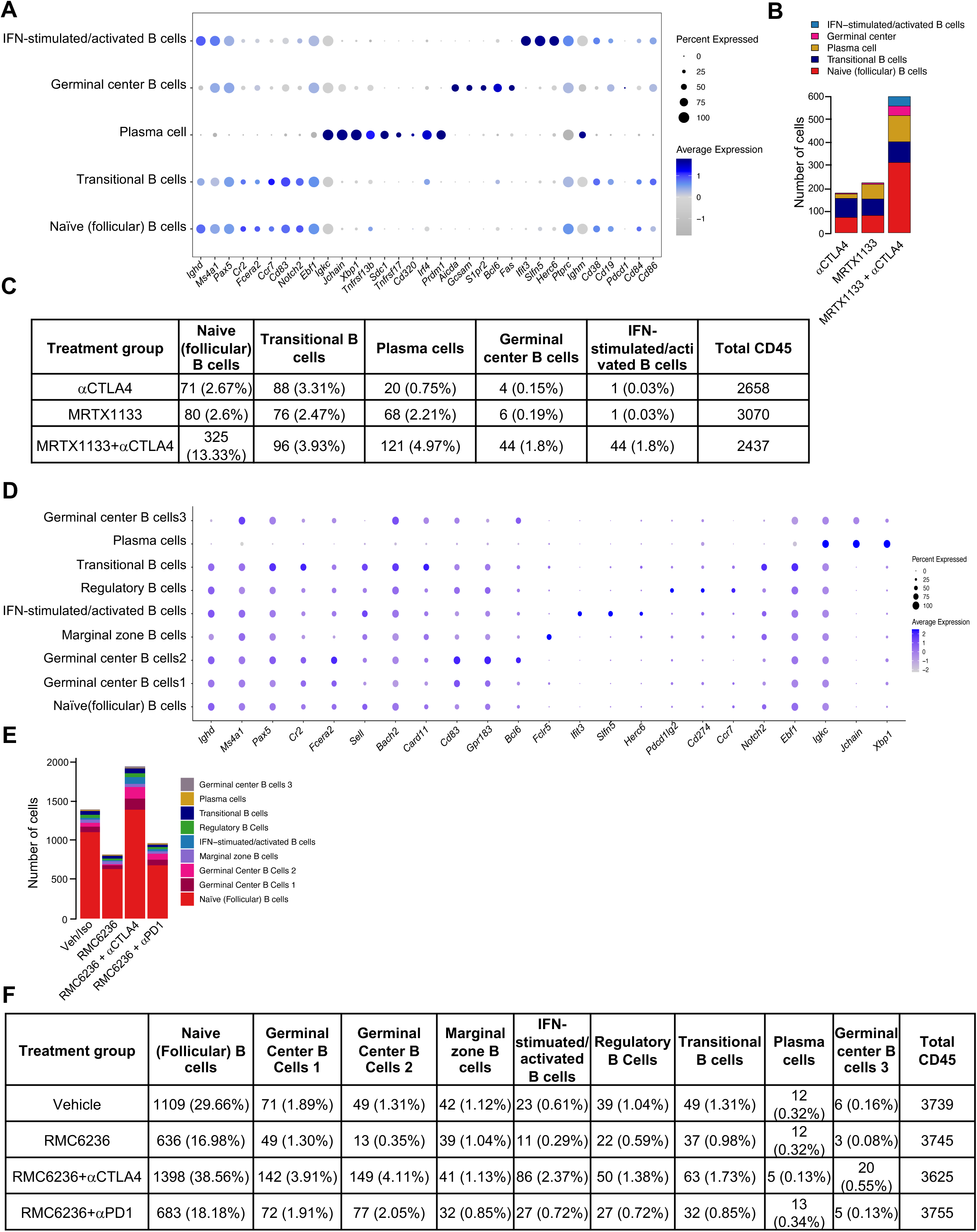
Additional scRNA seq. analysis of MRTX1133, RMC6236 treated PDAC. Related to Fig. 6. **(A)** Dot plots of genes used to define B cell subtypes in MRTX1133 treated KPC3 tumors determined by scRNA seq. analysis. **(B-C)** Number of B/ plasma cells in KPC3 PDAC as bar-graph **(B)** and table **(C)** determined by scRNA seq. from Fig. 6M-N. **(D)** Dot plots of genes used to define B cell subtypes in RMC6236 treated KPC2 tumors determined by scRNA seq. analysis. **(E-F)** Number of B/ plasma cells as bar-graph **(E)** and table **(F)** in KPC2 PDAC determined by scRNA seq. from Fig. 6O-P.

**Supplementary Fig. 16:**
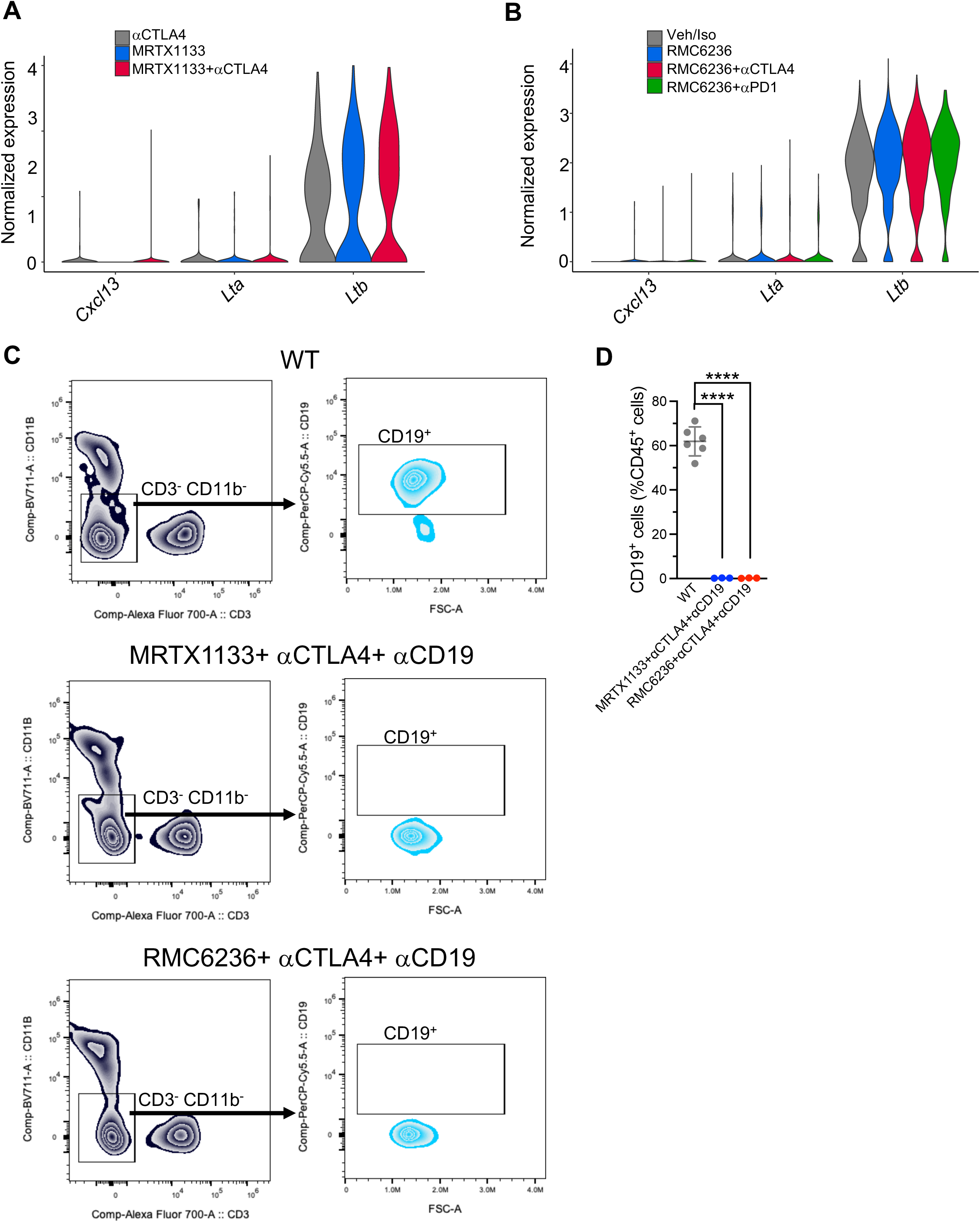
Additional analysis of scRNA seq. data and flowcytometry analysis. Related to Fig. 7. **(A)** Violin plots for *Cxcl13, Lta* and *Ltb* in tumor infiltrating B cells of MRTX1133 treated KPC3 mice with αCTLA4. **(B)** Violin plots for *Cxcl13, Lta* and *Ltb* in tumor infiltrating B cells of RMC6236 treated KPC2 mice with aCTLA4/aPD1. **(C-D)** Representative images of gating strategy in the peripheral blood of mice treated with MRTX1133/ RMC6236, aCTLA4 and aCD19 depletion antibodies **(C)**, with quantification of frequencies of indicated cell populations **(D)** (n=3-6 mice/ group). Data are presented as violin plots in **A, B** and as mean ± SD in **D**. Significance was determined by one-way ANOVA with Dunnett’s multiple comparisons test in **D**.

## Notes

### Summary of Updates

We have included GEM models and demonstrate synergy of Kras* inhibitors and synergy with anti-CTLA4 with accompanying tertiary lymphoid structures. Further, we demonstrate that the tertiary lymphoid structures are of functionally critical in mediating synergy of Kras* inhibitors with anti-CTLA4.

